# Dietary fibre deprivation and bacterial curli exposure shift gut microbiome and exacerbate Parkinson’s disease-like pathologies in an alpha-synuclein-overexpressing mouse

**DOI:** 10.1101/2022.03.21.485143

**Authors:** Kristopher J Schmit, Alessia Sciortino, Velma TE Aho, Pierre Garcia, Beatriz Pardo Rodriguez, Mélanie H Thomas, Jean-Jacques Gérardy, Rashi Halder, Camille Cialini, Tony Heurtaux, Irati Bastero Acha, Irina Ostahi, Eric C Martens, Michel Mittelbronn, Manuel Buttini, Paul Wilmes

## Abstract

The microbiome-gut-brain axis has been proposed as a pathogenic path in Parkinson’s disease (PD). Dietary driven dysbiosis and reduced gut barrier function could facilitate the interaction of toxic external or internal factors with the enteric nervous system, where PD could start. Amyloid bacterial protein such as curli can act as seed to corrupt enteric α-synuclein and lead to its aggregation. Misfolded α-synuclein can propagate to and throughout the brain. Here, we aimed at understanding if fibre deprivation and amyloidogenic protein curli could, individually or together, exacerbate the phenotype in both enteric and central nervous systems of a transgenic mouse overexpressing wild-type human α-synuclein. We analysed the gut microbiome, motor behaviour, gastrointestinal and brain pathologies in these mice. Our findings show that external interventions, akin to unhealthy life habits in humans, can exacerbate PD-like pathologies in mice. We believe that our results shed light on how lifestyle affects PD progression.

## Introduction

Lifestyle and environmental factors contribute to a variety of chronic, degenerative diseases burdening socio-economic structures in an expanding and ever-ageing population (Nations n.d.; Council (US) et al. 2013). The incidence rate of Parkinson’s disease (PD), the second most common neurodegenerative disease, has consistently increased over the last three decades (Dorsey et al. 2018) and is predicted to further increase (Yang et al. 2020). Of all current cases, only 5-10% can be attributed to heritable genetic factors alone (Poewe et al. 2017). For most cases, PD is a complex multi-factorial disease with genetic and environmental/lifestyle risk factors contributing to its onset and progression (reviewed in (Gorell et al. 2004)).

Environmental and lifestyle factors associated with PD are for example exposure to different chemicals (e.g. pesticides), head trauma, physical activity, stress, smoking and diet. Except for smoking and caffeine consumption, all other factors have been associated with increased risk of PD (reviewed in (Marras, Canning, and Goldman 2019)). In recent years, a growing body of evidence has put forward the importance of diet and its implications in PD progression. It has been proposed that a Mediterranean diet rich in fresh unprocessed foods, especially vegetables, reduces the risk for PD (Maraki et al. 2019). On the other hand, a “western” diet with low amounts of fibre and high amounts of saturated fats and simple carbohydrates has been associated, amongst a variety of other diseases, with neurodegenerative diseases such as PD (reviewed in (Hirschberg et al. 2019; Martínez Leo and Segura Campos 2020)and is a strong modulator of the gut microbiome. Functional comparative metagenomics analysis has shown that such a diet is associated with reduced gene expression related to complex carbohydrate fermentation (Rampelli et al. 2015). In different mouse models, low to fibre-free diets led to lower abundance of fibre fermenting bacteria (Schroeder et al. 2018) and higher abundance of mucus foraging species like *Akkermansia muciniphila* (Desai et al. 2016).

Desai and colleagues, amongst others (Martens, Chiang, and Gordon 2008), showed that this consequently led to increased mucus erosion and susceptibility to pathogens (Desai et al. 2016). While such pathogens rather come from infection (Nerius, Doblhammer, and Tamgüney 2020), there is evidence that such pathogens can also originate from commensal bacteria of the resident gut microbiome and contribute to the disease phenotype (Miller et al. 2021). Commensal bacteria occupy the outer mucus layer of the colon and can form biofilms (Johansson, Larsson, and Hansson 2011; Johansson, Sjövall, and Hansson 2013). Even though there is no consensus on biofilm formation in the healthy gut (Tytgat et al. 2019; De Vos and M 2015), it has been observed in a variety of gastrointestinal disease scenarios. One biofilm forming microbial family, *Enterobacteriaceae*, has been associated with severity of a specific subtype of PD (Scheperjans et al. 2015). Amongst the most prominent biofilm forming species are *Escherichia coli* (*E. coli*) and *Salmonella* (Miller et al. 2021). Both express curli, a major biofilm component(Miller et al. 2021) and amyloidogenic protein, which has structural and physiological similarities to ß-amyloid and α-synuclein (αSyn) (Chapman et al. 2002; Barnhart and Chapman 2006). It has been shown to act as a seed for αSyn aggregation *in vitro*(Sampson et al. 2020). When injected intramuscularly into the intestinal wall, αSyn aggregation was accelerated and led to motor deficits and GI dysfunction (Sampson et al. 2020). Further, colonization with curli expressing *E. coli* by oral gavage of young germ-free human wild-type αSyn overexpressing mice (Sampson et al. 2020) and microbiota-depleted Fischer 344 rats (S. G. Chen et al. 2016) did also replicate different aspects of PD including neuroinflammation, exacerbated motor deficits, and abnormal αSyn accumulations in gut and brain.

Alpha-synuclein accumulations in the gut have been observed in many PD patients (Wakabayashi et al. 1988; 1993; 1990; Qualman et al. 1984). Braak and colleagues proposed that αSyn accumulations in the ENS preceded those in the lower brainstem regions of the CNS and subsequently αSyn would propagate retrogradely in a “prion-like” manner via the vagus nerve to the brain (Braak et al. 2006). Interestingly, truncal vagotomy, the cutting of the vagus nerve near the gastroesophageal junction, has been associated with reduced risk for PD (Svensson et al. 2015; Liu et al. 2017). Under normal conditions, however, αSyn would further spread to the dorsal motor nucleus of the vagus and then follow Braak’s proposed trajectory leading to the common pathological hallmarks of abnormal accumulation of αSyn (Lewy bodies) and loss of dopaminergic neurons in the Substantia Nigra pars compacta (SNpc) and its projections in the dorsal striatum.

In this study we wanted to investigate the exacerbating effect of dietary fibre deprivation and bacterial curli exposure, individually or combined, on a human αSyn overexpressing transgenic mouse. Our subsequent treatment strategy, was to first prime the naïve untreated microbiome with a “westernized” fibre deprived diet (Desai et al. 2016), followed by the exposure to curli producing bacteria (S. G. Chen et al. 2016). We analysed the mice at gut microbial, behavioural, gastrointestinal, and neuropathological levels. Overall, transgenic, but not wild-type mice, were susceptible to the different challenges. In transgenic mice, our findings suggest that even though αSyn overexpression is greatly responsible for the observed behavioural impairments, the fibre deprived caused dysbiosis led to increased mucus erosion and pathogen susceptibility, consequently resulting in PD-like pathologies, such as αSyn accumulation in the enteric nervous system (ENS) and nigro-striatal degeneration and αSyn accumulations in the central nervous system (CNS) further exacerbating coordinative skills of our transgenic mice. We believe that our study sheds light on how a combination of internal and external pathogenic factors can differentially contribute to PD-like pathologies in the CNS and ENS. Therefore, our findings may have important implications for lifestyle adjustments that could mitigate PD.

## Results

### Thy1-Syn14 overexpress alpha-synuclein in brain and gut with regional differences

The mouse model used in this study was first mentioned in (P. J. Kahle et al. 2001). It overexpresses human wild-type αSyn under the transcriptional regulation of the neuron specific Thy1 promotor and carries 13 copies of the transgene (P. J. Kahle et al. 2001). The model has so far not been fully described in literature, and only protein levels in bulk brain tissue have been reported (P. J. Kahle et al. 2001). Thus, determining baseline expression and protein levels of αSyn in different CNS regions and in the gut was crucial for the subsequent interpretation of data generated in our study.

In the CNS, we focused on ventral midbrain and dorsal striatal structures using RT-qPCR for gene expression and Western blot for protein level quantifications. We focused on both the differences between genotypes and the regions. First, we saw that gene expression levels for the murine αSyn (*Snca)* gene (**Fig. 1a**, top row) did no change between genotypes, but were significantly different (p < 0.0001) between the ventral midbrain and dorsal striatum. On the other hand, for the human transgene of αSyn (**Fig. 1a**, bottom row), we could only detect signal in transgenic mice and those levels did not differ significantly between regions. We saw similar protein profile changes for total αSyn protein levels. To measure total αSyn protein levels, we used a pan-αSyn antibody. This antibody detects both murine and human αSyn. We observed significant differences between genotype for each region and between regions (WT: p = 1.52E-2; TG: p = 1.55E-4; **Fig. 1b**). For transgenic mice, we measured a 2.91-fold increase (p = 6.67E-4) in the dorsal striatum and a 6.67-fold increase (p = 6.67E-4) in the ventral midbrain compared to their wild-type littermates (**Fig. 1b**). The observed distribution profile showed us that the expression of murine αSyn (Snca, **Fig. 1c**, right column) varied between the regions of interest, no matter the genotype, while human αSyn (SNCA, **Fig. 1c**, middle column), aside from only present TG mice, was expressed homogeneously in all regions. Finally, the total αSyn expression profile was identical to what we observed for SNCA. This reflected the high protein level differences observed by Western blot.

**Fig. 1.**
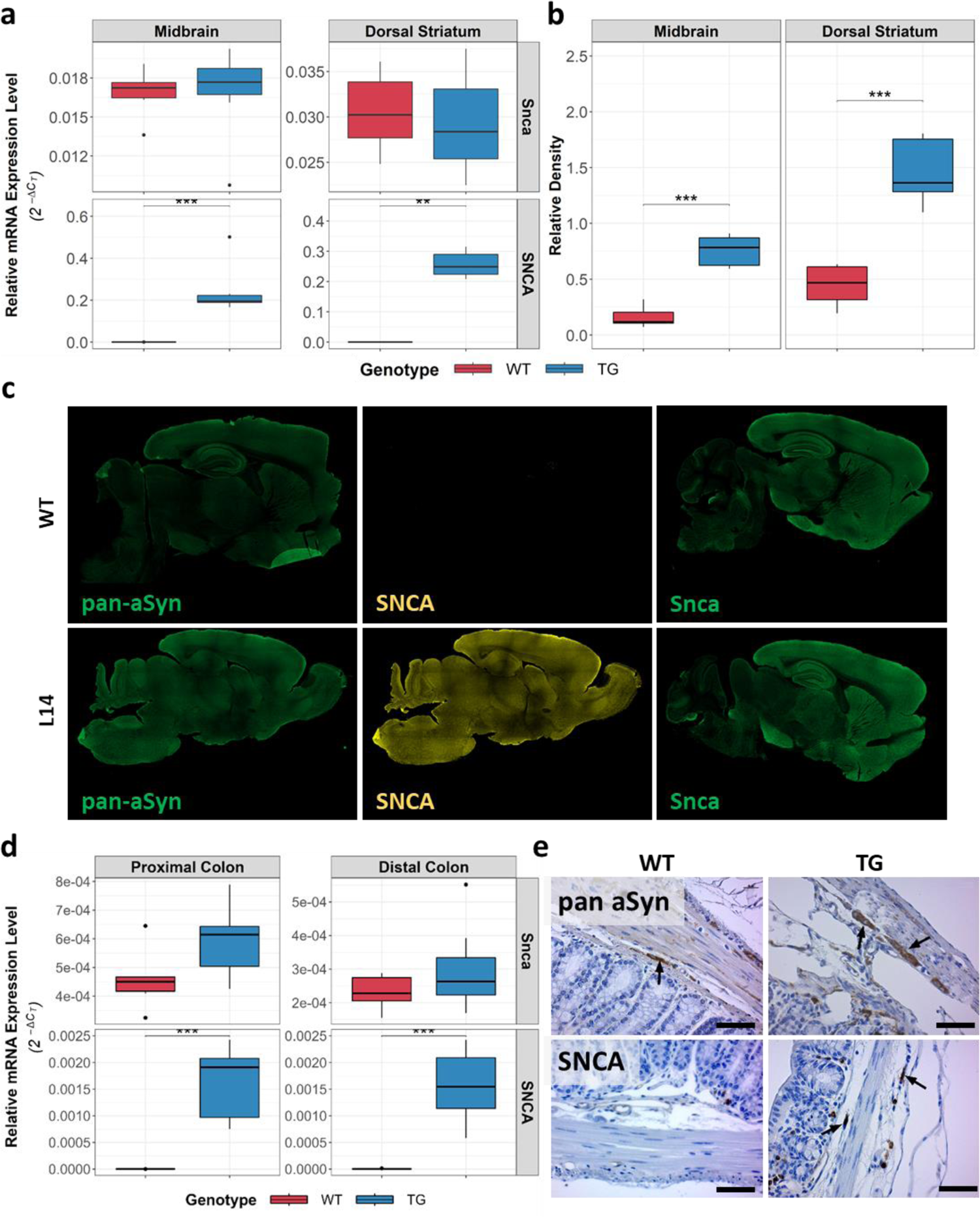
Thy1-Syn14 present high alpha-synuclein gene expression and protein levels with regional differences. a) Boxplots showing relative mRNA expression levels of Snca (bottom panel) and SNCA (top panel) of 9 months old Thy1-Syn14 (TG, blue) mice and wild-type littermates (WT, red) in midbrain and dorsal striatum. Only TG mice express human αSyn. There were no differences for Snca between WT and TG animals per region. b) Boxplots illustrating the calculated relative density from Western blots for pan-αSyn in the ventral midbrain and dorsal striatum. Alpha-tubulin was used as reference protein. c) Representative immunofluorescent stainings for pan-αSyn, mouse αSyn (Snca) and human αSyn (SNCA). Human αSyn is absent in WT and homogeneously expressed in TG animals (middle panel). Murine αSyn, is expressed uniformly, except for the ventral midbrain, and there are no differences between genotypes (right panel). Qualitatively, pan-αSyn is more readily expressed in TG mice. d) Boxplots showing relative mRNA expression levels of Snca (top panel) and SNCA (bottom panel) for 9 months old TG (blue) and WT (red) mice in proximal and distal colon. Only TG animals express human αSyn (SNCA) without regional differences. Murine αSyn appears to be higher expressed in proximal colon samples and more so in TG animals. e) Representative images (Scale bar: 50µm) from immunohistochemistry stainings showing that SNCA in enteric neurons (arrow, right lower panel) is only expressed in TG animals, while both WT and TG are positive for pan-αSyn (top row panel, black arrows). Stats: Kruskal-Wallis test; **, p<0.01; ****, p<0.001 WT, wild-type littermates, TG, Thy1-Syn14 animals, pan-αSyn, total alpha-synuclein protein

Next, we checked the αSyn expression profile in the colon. We split the colon into proximal and distal parts and measured both mouse (*Snca*) and human (SNCA) αSyn expression levels via RT-qPCR. It was apparent that overall the levels for *Snca* and *SNCA* are much lower when comparing to the observed CNS levels (**Fig. 1d**). This is most likely due to the much lower density of neurons in whole colon compared to e.g. ventral midbrain samples. In colon, only about 1% of all cells are neurons (Drokhlyansky et al. 2020), whereas e.g. ventral structures of the midbrain, comprising several populations of dopaminergic neurons (e.g. Substantia Nigra and Ventral Tegmental Area), have a roughly estimated proportion of 15-20% neurons (Keller, Erö, and Markram 2018; Murakami et al. 2018; J. Zhang et al. 2007; Y. Zhang et al. 2012). Nevertheless, we saw that *SNCA* was only expressed in transgenic mice, while there were regional but no genotype differences for *Snca*. Additionally we stained for human αSyn and total αSyn (**Fig. 1e**). Latter was expressed in both genotypes, while only transgenic mice expressed human αSyn (**Fig. 1e**, left bottom panel). The data shown here, indicates that the Thy1-Syn14 model presents an appropriate system to investigate αSyn related gut-brain interactions.

### Thy1-Syn14 mice exhibit progressive motor deficits

PD-like motor deficits have been reported in a variety of mouse models of the disease (reviewed in (Barber Janer, Vonck, and Baekelandt 2021)). Here we assessed for grip strength, hind limb reflexes and coordination/movement.

To assess grip strength, indicative of striatal dysfunction and neurotransmitter loss (Tillerson et al. 2002a), we used a simplified version of the inverted grid test (Tillerson et al. 2002b; Tillerson and Miller 2003). Here, we only measured the hanging time and therefore assessed the simultaneous 4-limb grip strength. Grip strength gradually decreased as the TG mice aged (3M: 16.2 ± 9.88s, p = 3E-4; 6M: 11 ± 9.39s, p = 3E-4; 13M: 0.17 ± 0.41s, p = 4.86E-7; **Fig. 2a**). Grip strength in wild-type littermates remained unchanged.

**Fig. 2.**
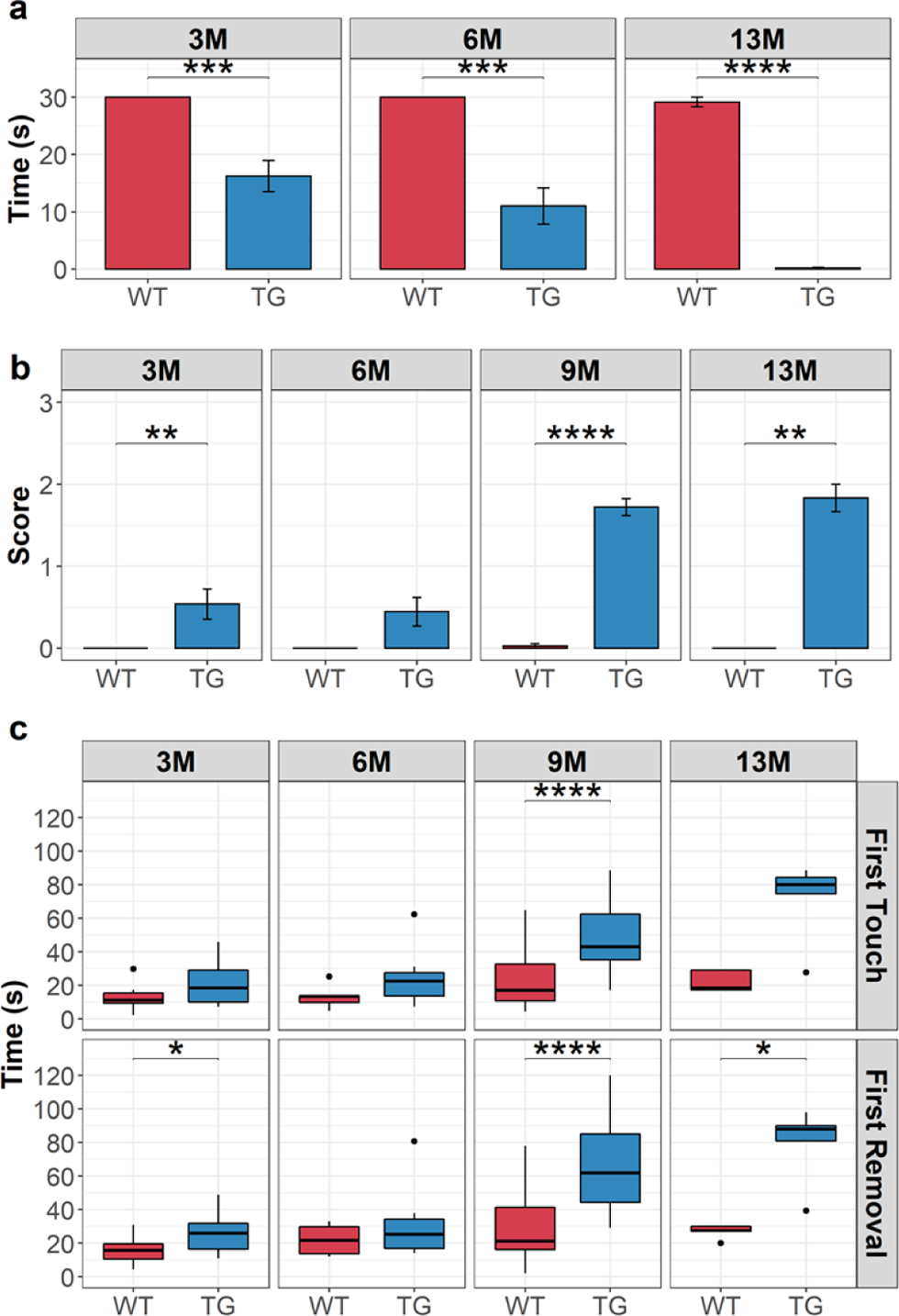
Thy1-Syn14 showed a progressive decline in behavioral motor skills. a-c) Animals were tested at different ages (3M, 6M, 9M, 13M) for gross and fine motor skills. Results are depicted in boxplots indicating median values with interquartile ranges. a) Inverted grid indicates that animals gradually lose grip strength. Muscle weakening is already significantly reduced at three months of age. At 6 months of age, grip strength declines further, and at 13 months of age, transgenic animals cannot hold on to the grid anymore at all. Data for 9 month old animals is missing. b) Hind-limb clasping appears also as early as three months in some animals. When specifically looking into age dependant responses, animals slightly older than three month were more likely to show single hind-limb clasping (score = 1). In the three and 6 month cohorts there was no clasping of both hind-paws (score = 2). Only at 9 months and to a greater extend at 13 months animals showed clasping of both hind-paws. Wild-type animals did not show (with one exception at 9 months) hind-limb clasping at any age on average. c) Adhesive removal measure for fine sensory and motor coordination skills. Results indicated that these skills are age dependant and differed significantly between wild-type and transgenic animals in the 9 and 13 months cohorts. Stats: Mann-Whitney U, corrected for FDR; *, p < 0.05, **, p < 0.01, ***, p < 0.001, ****, p < 0.0001 WT, wild-type littermates; TG, Thy1-Syn14 animals, 3M, 3 months old; 6M, 6 months old; 9M, 9 months old; 13M, 13 months old

The hindlimb clasping or reflex test, an additional test assessing striatal dysfunction (Fernagut et al. 2004; Lieu et al. 2013), confirmed that the performance of TG mice decreased with age (3M: p = 9.33E-3; 6M: p = 5.7E-2; 9M: p < 0.0001; 13M: p = 4E-3; **Fig. 2b**), while we did not observe an age-dependent motor impairment in the wild-type littermates.

Finally, we tested mice for coordination and fine motor skills with the adhesive removal test. Deficits in time of removal have been associated with loss of dopaminergic neurons (Fleming, Ekhator, and Ghisays 2013). We did not observe relevant motor deficits in young mice (3M and 6M; **Fig. 2c**). At 9 and 13 months, we saw that both sensitivity (Time at touch, upper strip), as well as coordination (Time at removal, lower strip) were significantly delayed in TG mice (Time at touch – 9M: p = 47.57E-7, 13M: p = 0.057; Time at Removal – 3M: p= 0.016, 9M: p = 1.12E-6, 13M: p = 0.016). Taken together, these data indicate that overexpression of human wild-type αSyn drives progressive motor dysfunction in the Thy1-Syn14 mice.

### Translationally relevant PD-like microbial changes induced after fibre deprivation

Onset and progression of PD, especially idiopathic PD, has been linked to the exposure of different environmental factors (H. Chen and Ritz, n.d.; Di Monte, Lavasani, and Manning-Bog 2002; Dick et al. 2007; Warner and Schapira 2003), of which some, e.g. diet, impact the gut microbiome(Bernardo-Cravo et al. 2020; Singh et al. 2019). Changes in gut microbial composition in PD has been described extensively in humans(Boertien et al. 2019; Gerhardt and Mohajeri 2018; Heintz-Buschart et al. 2018; Keshavarzian et al. 2015; Scheperjans et al. 2015; Shen et al. 2021; Unger et al. 2016), but also more and more in different animal models(Gorecki et al. 2019; Sampson et al. 2016; Yan et al. 2021). Using 16S rRNA amplicon sequencing, our goal was to understand how the disease challenges affected, independently or in combination, the microbial phenotype in our mice.

First, we looked on a large how the challenges affected microbial diversity. Overall, we identified the FD diet and TG challenges to be implicated in reduced inner-group diversity (alpha diversity), while no changes were observed for the gavage challenges (**Supplementary Fig. 2**). Alpha diversity was lowest at weeks 2 (WT: p = 0.045; TG: p = 6.1E-4) and 9 (TG: p = 9.7E-3) in FD challenged and more prominently so in TG mice (**Fig. 3a**). We made similar observations for beta diversity, where the diet challenge was the main driver (adonis p = 0.001) of dissimilarities (**Supplementary Fig. 2a**, middle panel). However, different to alpha diversity, the gavage challenges (adonis p = 0.001) appeared to contribute as well (**Supplementary Fig. 2b**, right panel). This observation might however be due to the PBS gavaged mice having all been FD challenged. Hence, solely the FD challenge drove the observed rapid microbial shift (**Fig. 3b**). Already at week 2, the FD and FR challenged groups, independent of the other challenges, formed two homogeneous clustered until the end of the experimental in-life phase. Such a shift is indicative of dysbiosis, the functional imbalance of the microbiome. Changes in the *Firmicutes* to *Bacteriodetes* ratio, the two most abundant phyla in the gut, can hint to an overall microbial and functional imbalance (Magne et al. 2020; Mariat et al. 2009). Our data showed significant increases in the *Firmicutes* to *Bacteriodetes* ratio in FD challenged mice (**Supplementary Fig. 3**). Next, we focused our analysis on taxa abundance changes at the genus level.

**Fig. 3.**
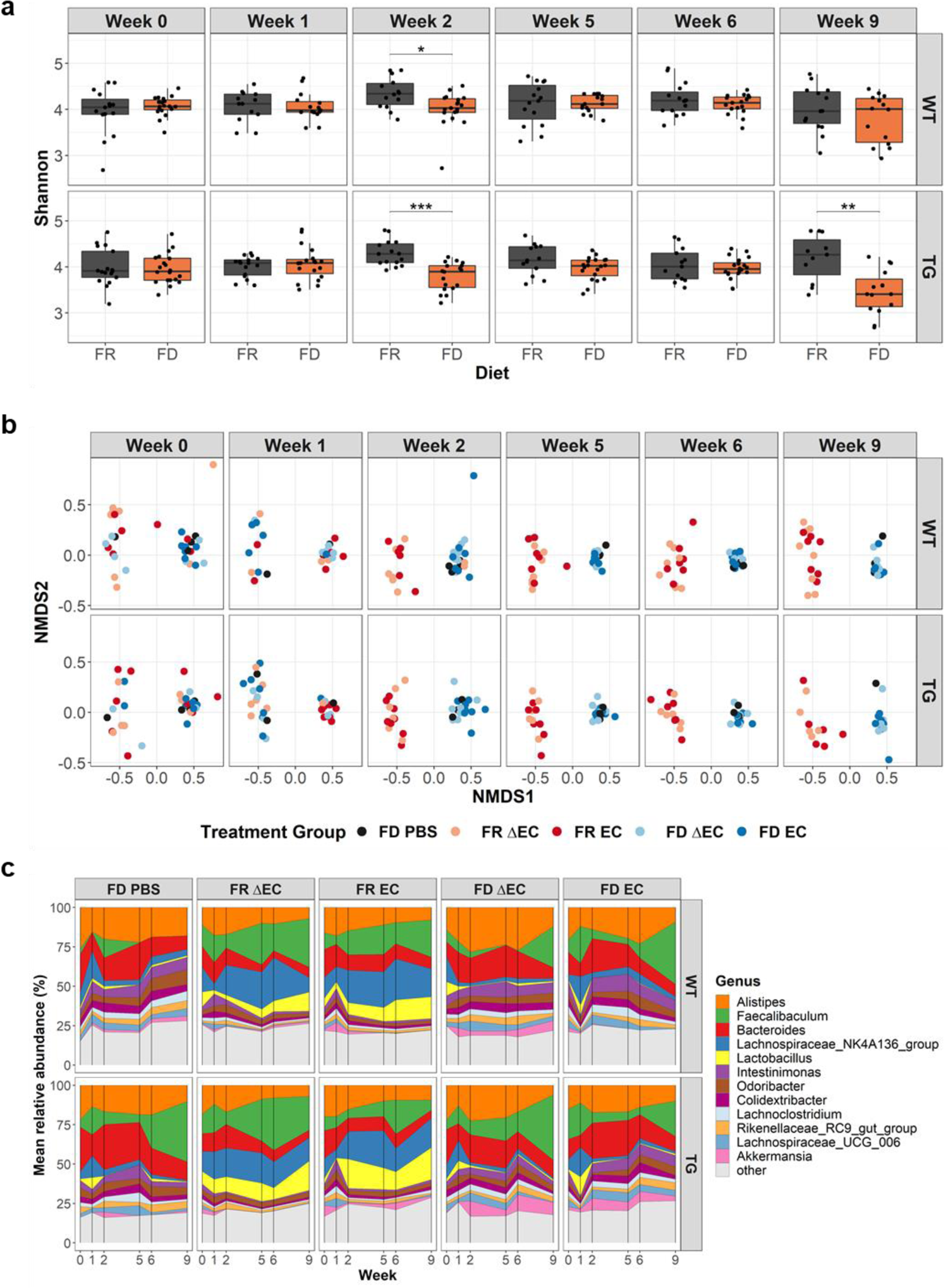
Longitudinal changes in microbial diversity and composition shifts are dietary driven. a) Boxplots illustrating alpha diversity for the two diet challenges at different time points and both genotypes separately. The absence of dietary fibre (FD) reduced microbial diversity significantly at week 2 in WT (FDR < 0.05) and TG (FDR < 0.001) animals. At week 2 we started the gavages. In TG animals microbial diversity is again significantly (p < 0.01) reduced at week 9. Stats: Mann-Whitney U corrected for FDR; *, p < 0.05; **, p < 0.01; ***, p < 0.001. b) Non-metric multi-dimensional scaling (NMDS) representations for beta diversity showing the different treatment groups facetted by genotype (row) and the different time points (Week, column). We observe a composition shift leading to two homogeneous clusters by week 2. This separation is driven by the dietary challenges. c) Temporal distribution of relative abundance for the 12 most abundant and relevant taxa on the genus level for weeks 0 (Baseline), 1, 2, 5, 6 and 9. Both WT (top row) and TG (bottom row) mice show similar changes in relative abundance for the different taxa. Differences were observed between the diet challenge groups. WT, wild-type littermates; TG, Thy1-Syn14 mice; FD, fibre deprived; FR, fibre rich; PBS, phosphate buffered saline solution; ΔEC, curli-KO E.coli; EC, wild-type curli expressing E.coli

We first subdivided the taxa according to their abundance into three separate groups (high, mid, and low; **Supplementary Fig. 4**). Analogous to what has been seen before, the diet challenge was the main driver of microbial abundance changes (**Supplementary Fig. 4**). Next, we focused our analysis on the most relevant genera in our data, which were *Alistipes, Faecalibaculum, Bacteroides, Lachnospiraceae NK4A136 group, Lactobacillus, Intestinimonas, Odoribacter, Colidextribacter, Lachnoclostridium, Lachnospiraceae UCG 006, Rikenellaceae RC9 gut group* and *Akkermansia* (**Fig. 3c**). FD challenged mice had increasing or constantly higher levels in *Faecalibaculum, Intestinimonas, Odoribacter, Colidextribacter, Rikenellaceae RC9 gut group* and *Akkermansia,* and decreasing levels in *Lachnospiraceae NK4A136 group* and *Lactobacillus* over time (**Fig. 3c, Supplementary Fig. 4**). *Alistipes, Bacteroides, Lachnoclostridium* and *Lachnospiraceae UCG_006* on the other hand saw fluctuations over time. They first increased in abundance before dropping back to initial levels in FD challenged mice.

Next, when compared to data from PD patients (Table 3 in Boertien et al., 2019(Boertien et al. 2019)), FD challenged mice showed similar changes in *Akkermansia* (and its corresponding Family and Phylum), *Lachnospiraceae*, *Roseburia* and *Prevotellaceae*, while *Lactobacillaceae* and its genus *Lactobacillus* were inversely altered (**Supplementary Table 1**). Other taxa that are often reported to be dysregulated in human PD stool samples were not detected, e.g. *Bifidobactericaceae, Faecalibacterium* (*Clostridiaceae*) or *Enterobacteriaceae* (**Supplementary Table 1**). When we checked for the relative abundance of *Escherichia coli*, the *Enterobacteriaceae* species that we gavaged, it was barely detected in our 16S rRNA amplicon sequencing data (**Supplementary Fig. 5**).

In summary, the FD challenge caused reduced gut microbial diversity and similar shifts as seen in PD patients. To note are reduced levels of *Lactobacillus*, a known probiotic(Heeney, Gareau, and Marco 2018; Martín et al. 2013) with possible neuroprotective effects(Wang et al. 2021) and associated with gut barrier integrity(Blackwood et al. 2017), and *Lachnospiraceae NK4A136 group*, which inversely correlates with risk for PD or dementia(Stadlbauer et al. 2020). Additionally, *Lachnospiraceae NK4A136 group* and *Roseburia* are important butyrate producers, thus associated with gut barrier function(Plöger et al. 2012). Consequently, even though plasma endotoxin levels were not significantly increased in FD challenged mice (**Supplementary Fig. 6**), it is reasonable to assume that FD challenged mice had reduced gut barrier function. Together with higher levels of the known mucin-foraging genera *Akkermansia* and *Bacteroides*(Desai et al. 2016; Tailford et al. 2015), the susceptibility for pathogenic factors was most likely increased.

### Microbial driven mucus erosion of the colon in fibre deprived challenged mice

The mucus layers of the colon have two basic functions: the inner layer acts as physical barrier to prevent pathogens of reaching the gut epithelium and the outer layer harbours commensal bacteria interacting with the host(Johansson, Sjövall, and Hansson 2013). Increasing levels of mucin degrading bacteria, such as *Akkermansia muciniphila* and certain *Bacteroides* spp.(Desai et al. 2016), have been shown to cause mucus thinning consequently facilitating epithelial access for potential pathogens.

For this study we focused on the outer mucus layer. Our results clearly showed that the FD challenge caused significant (p = 1.15E-12) thinning of the outer mucus layer (**Fig. 4a**). The layer thickness decreased extensively by 49.1% to 92.9% due to the diet challenge (**Fig. 4b**). Hence, the habitat for gut bacteria was reduced having direct consequences on microbial diversity. Therefore, we compared alpha diversity and mucus thickness using the Spearman’s rank test. We were specifically interested in dietary or transgene driven associations. While we only saw moderate dietary driven correlations (FR: r=-0.47, p=0.051; FD: r=0.38, p=0.08; **Supplementary Fig. 7**), there was a significant positive correlation between alpha diversity and mucus thickness in TG mice (**Fig. 4c**) pointing to an αSyn-related increased susceptibility to microbial changes.

**Fig. 4.**
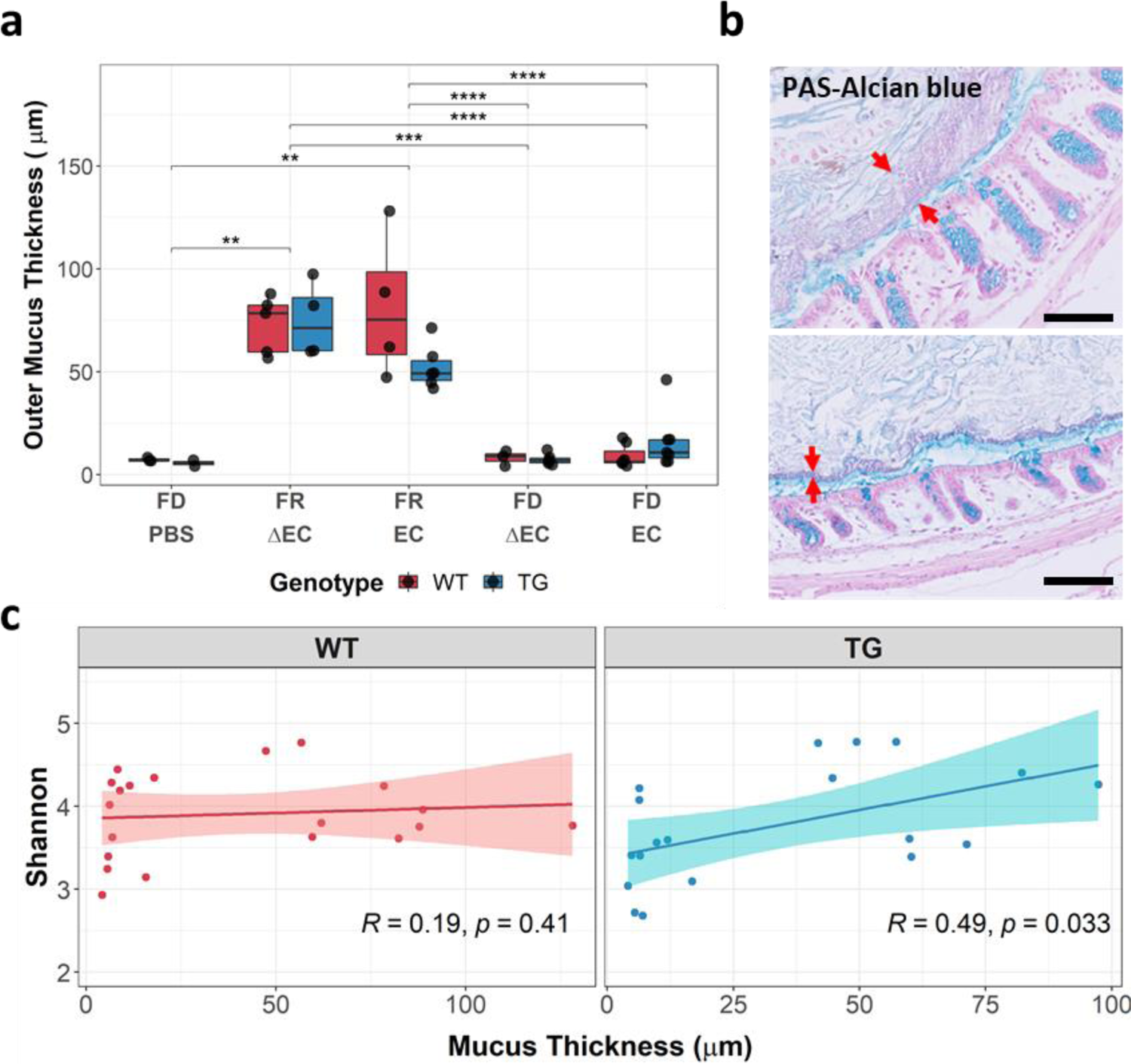
Microcbial mediated outer mucus erosion associated to reduced diversity. a) Boxplot of outer mucus thickness measurements. The FD challenge causes a vast reduction in outer mucus thickness. Stats: Mann-Whitney U test, corrected for FDR; **, p < 0.01; ***, p < 0.001; ****; p < 0.0001. b) Representative images illustrating the differences in outer mucus erosion between FR (top) and FD (bottom) challenged mice. The red arrows delimit the outer mucus layer. c) Scatterplots of Spearman rank tests comparing alpha diversity (y-axis) and mucus thickness (x-axis) in both genotypes separately. There is a significant positive correlation between microbial diversity and mucus thickness in Thy1-Syn14 animals independent of the other challenges. Stats: Spearman rank test. WT, wild-type littermates; TG, Thy1-Syn14; FD, fibre deprived; FR, fibre rich; PBS, phosphate buffered saline solution; ΔEC, curli-KO E.coli; EC, wild-type curli expressing E.coli

### Bacterial curli drives alpha-synuclein accumulation in the colonic myenteric plexus in fibre deprived challenged Thy1-Syn14 mice

Alpha-synuclein accumulation in the gut has been observed in a substantial number of PD patients(Braak et al. 2006; Del Tredici and Braak 2012; Del Tredici and Duda 2011). To test for increased levels of αSyn levels in the gut of our mice, we used an antibody directed against phosphorylated S129 αSyn (pS129-αSyn). This kind of antibody is commonly used to detect αSyn accumulation in both the murine and human central nervous system (CNS)(Vaikath et al. 2019) and enteric nervous system (ENS)(Shannon et al. 2012; Stokholm et al. 2016). We quantified the pS129-αSyn positive accumulations in protein gene product 9.5 (PGP9.5), a neuronal cytoplasmic marker(Sidebotham et al. 2001), positive ganglions of the myenteric (or Auerbach) plexus. Other pS129-αSyn positive accumulations in the in the submucosal plexus and submucosa were irregular and mainly detected in TG mice. This did however not differ between the different challenge group and we could not determine the cell type with our approach.

In our analysis, we focused on the myenteric plexus. First, the stainings showed that both WT and TG mice had pS129-αSyn positive accumulations in ganglions of the myenteric plexus (**Supplementary Fig. 8**). The quantification of the area occupied showed that only TG mice exposed to the combined challenge FD EC had increased levels of pS129-αSyn in PGP9.5 positive ganglions (vs TG FD PBS: p = 0.019; vs WT FR EC: p = 0.019; vs TG FR EC: p = 0.008; vs WT FD ΔEC: p = 0.019; **Fig. 5**). Besides having a greater area occupied, the average particle size of the αSyn accumulations appeared increased in TG FD EC challenged mice (representative images **Fig. 5**).

**Fig. 5.**
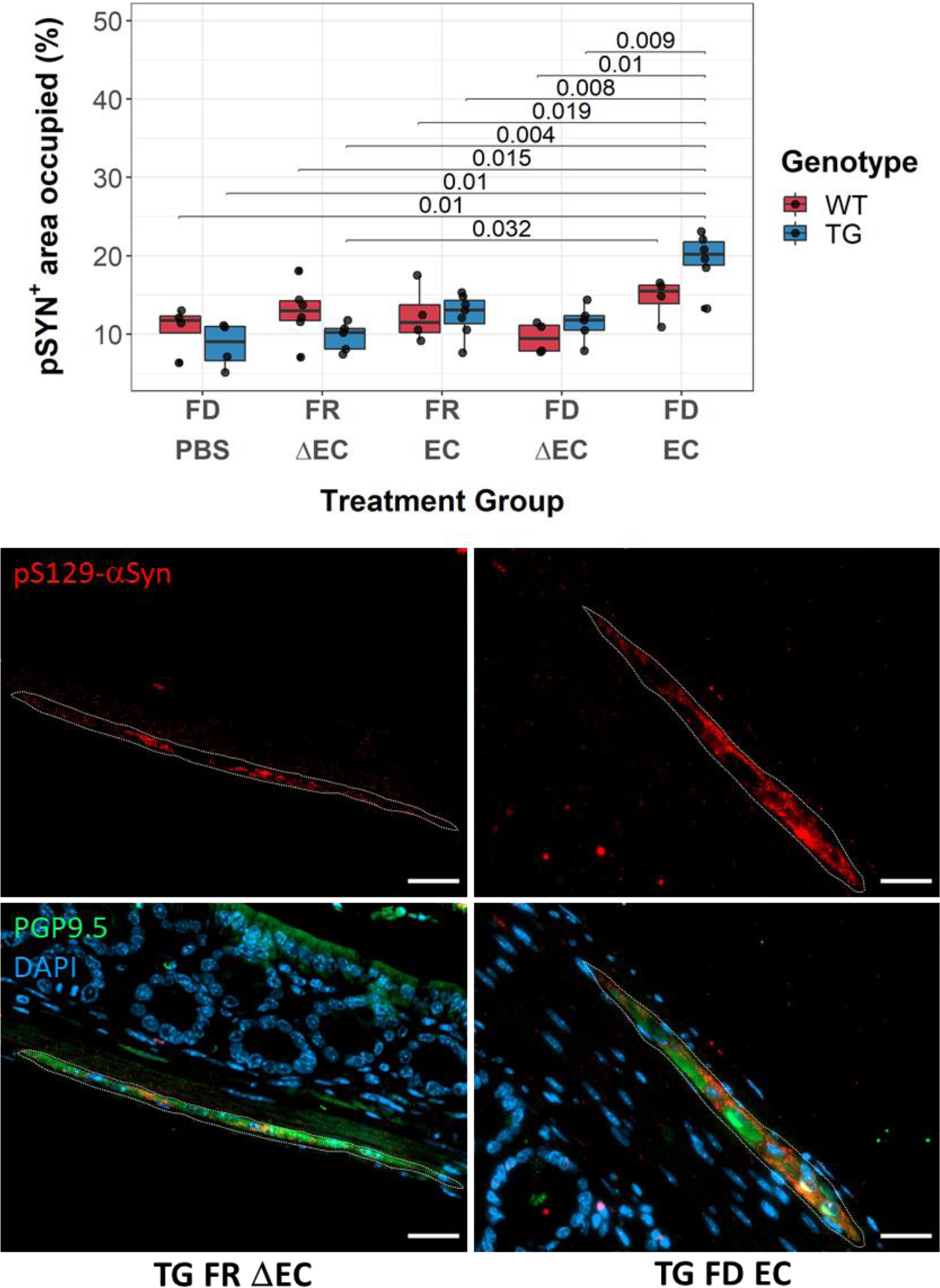
Curli-driven phospho-synuclein accumulation in fibre deprived challenged Thy1-Syn14 mice Boxplot illustrating the changes of area occupied by pS129-αSyn+ forms in ganglions of the myenteric plexus of the colon. Only TG mice on the combined challenge have an increased area occupied for pS129-αSyn+ forms. Stats: Mann-Whitney U test, not corrected for FDR. Representative images below, illustrate the average differences between FR ΔEC and FD EC challenged Thy1-Syn14 mice. Besides the area occupied, the pS129-αSyn+ particles are also enlarged in FD EC challenged mice. WT, wild-type littermates; TG, Thy1-Syn14; FD, fibre deprived; FR, fibre rich; PBS, phosphate buffered saline solution; ΔEC, curli-KO E.coli; EC, wild-type curli expressing E.coli

We can sum up that even though, as seen above, the FD challenge in general led to an increased pathogen/pathobiont susceptibility, only TG EC challenged mice showed significantly higher levels of pS129-αSyn positive accumulations in the myenteric plexus of the colon.

### Alpha-synuclein overexpression is the main but not sole driver of behavioural changes

The exposure to curli, here and in previous studies(S. G. Chen et al. 2016; Sampson et al. 2020), has been shown to lead to increased accumulation of abnormal αSyn in the gut. The subsequent spreading to and within the brain has been associated to motor impairment progression, as hypothesized by Braak and colleagues(Braak et al. 2003; 2006). In our aged TG mice, we did already observe motor performance deficits compared to their WT littermates. Therefore, we turned our interest to the exacerbation of motor performance deficits after the challenges, individually or in combination.

Three different tests were chosen to assess motor performance: hindlimb clasping, grip strength and adhesive removal. Hindlimb clasping and grip strength are so called basic or gross motor function tests and were used to monitor motor performance changes along the in-life phase of the study. Already at baseline, our aged mice differed significantly in their clasping and grip strength phenotype (**Supplementary Fig. 9a, 9b**). Over the course of the experiments only TG mice showed changes for both features (**Supplementary Fig. 9a, 9b**). The behavioural changes in TG mice were, however, neither linked to the diet, nor the gavage challenges. Hence, the progressive impairment of these features was driven by the overexpression of αSyn.

The adhesive removal test was used to assess changes in fine motor function (see Material and Methods). The test consists in measuring the latencies of touch and removal, which at baseline were significantly (p < 0.0001) greater in TG mice compared to their WT littermates (**Fig. 2c, 9M**). This was still the case at the end of the 9-week long experimental phase (p < 0.0001; **Fig. 6a, Supplementary Fig. 10**). Hence, αSyn overexpression was again key in driving motor impairment. For the external challenges, particularly TG EC challenged mice, independent of the diet, showed increased times of touch and removal, respectively, compared to the WT littermates (**Fig. 6a, Supplementary Fig. 10**). Further, we saw a much greater increase in both measures for a subset of TG FD EC challenged mice. This raised additional questions: how did the performance change over time for these TG mice? And what is the interval between the time of touch and time of removal? To answer these questions, we 1) subtracted the time of touch from the time of removal and then 2) compared “Baseline” to “Endpoint” results. We found that for the TG FD EC group 3 out of 6 mice had reduced coordinative ability to remove the adhesive tape compared to their initial performance (**Fig. 6b**). While statistically we did not get significant differences, the combination of the diet and curli challenges did appear to further exacerbate the motor phenotype in aged TG mice.

**Fig. 6.**
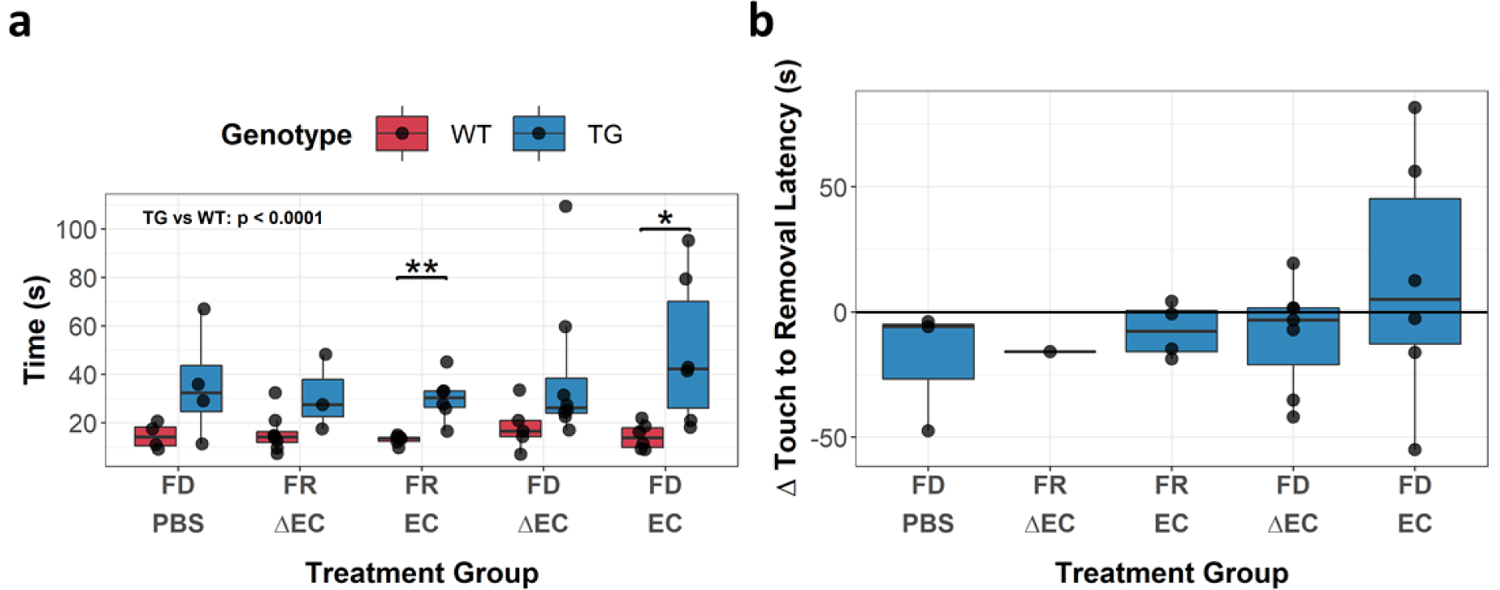
Challenge effect on motor impairment in a subset of Thy1-Syn14 mice despite the strong age-related transgenic phenotype. a) Boxplots illustrating the latency for removal in the different treatment groups (x-axis) for WT (red) and TG (blue) animals after 9 weeks. The results shown are for the first paw and from the first replicate. WT animals do not show any difference. TG mice on the other hand that have been FD EC challenges showed greater latency to remove the adhesive tape. There are significant in the FR EC and FD EC groups between WT and TG mice. Stats: Mann-Whitney U test, not corrected for FDR; *, p < 0.05; **, p < 0.01. b) Summary plot for TG animals illustrating how the adhesive removal performance changed from baseline to endpoint when focusing on the time difference from touch to removal. Most animals improved in performancefrom baseline to endpoint. Only the FD EC group shows for 3 out of 6 animals a performance drop. Note: for group 7 only one animal performed normally and so this group result can/should be neglected. (FD PBS: n=3, FR ΔEC: n=1, FR EC: n=5, FD ΔEC: n=6, FD EC: n=6). WT, wild-type littermates; TG, Thy1-Syn14; FD, fibre-deprived; FR, fibre-rich (normal chow); PBS, phosphate buffered saline; ΔEC, curli-KO E.coli; EC, wild-type curli expressing E.coli

### Bacterial curli mediated accumulation of alpha-synuclein in the nigrostriatal pathway

All neuropathological analyses were limited to TG mice, since they have shown to be more susceptible to the challenges. In a first step, we wanted to determine abnormal αSyn accumulations by immunofluorescence staining in the nigrostriatal pathway. To do so, we used again the pSy129-αSyn antibody and quantified pS129-αSyn positive accumulations in the dorsal striatum and the substantia nigra pars compacta (SNpc). Overall, we observed that the EC challenge was the main driver of pS129-αSyn accumulation in both regions of interest (**Fig. 7**).

**Fig. 7.**
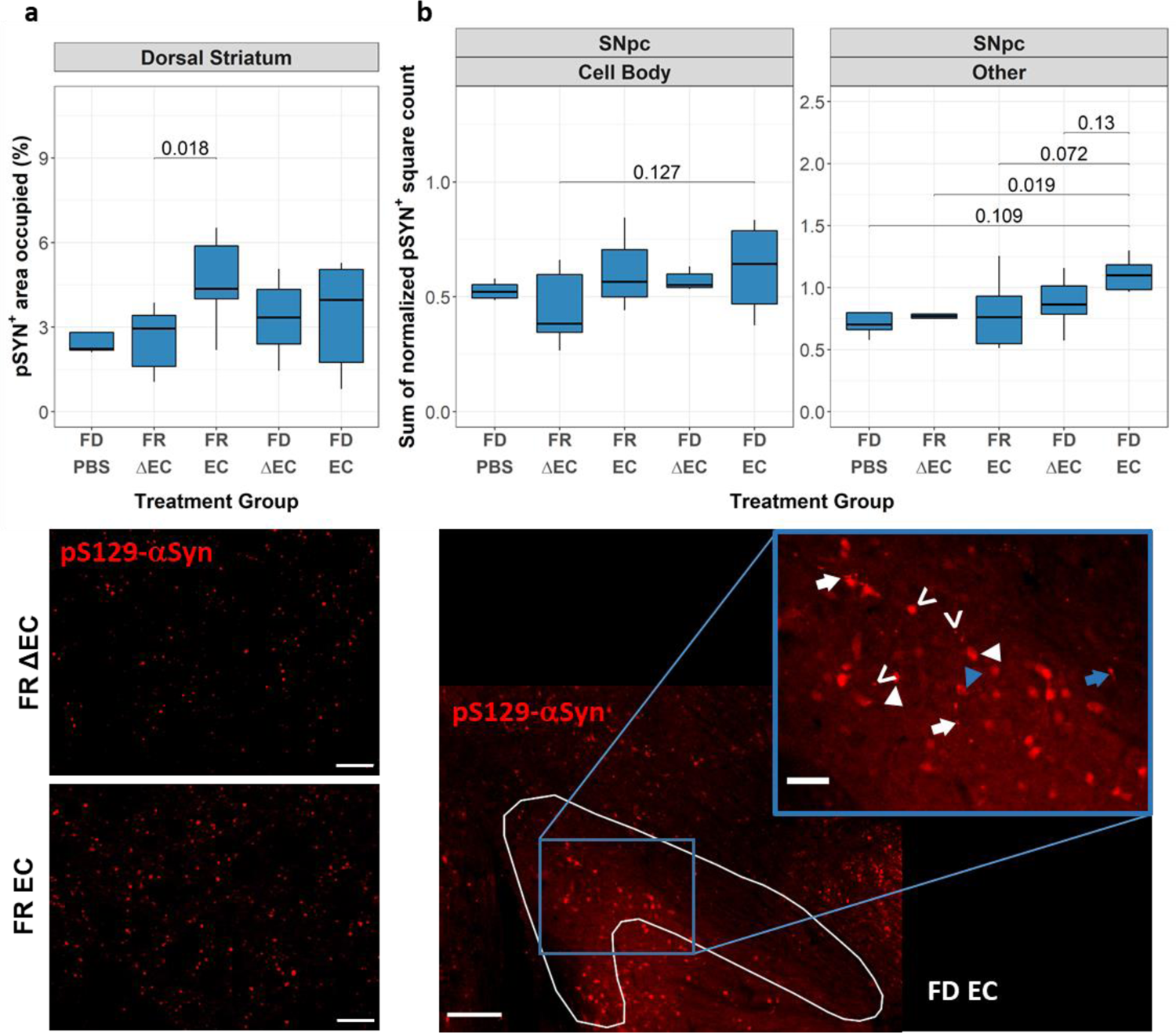
Curli-driven nigrostriatal phospho-synuclein accumulations in Thy1-Syn14 mice. a,b) Quantification and representative images of immunofluorescent pS129-αSyn stainings of the nigrostriatal pathway. a) Quantitive analysis of pS129-αSyn accumulations in the dorsal striatum by measuring the relative pS129-αSyn+ occupied area. The representative images (40X, scale bar: 20µm) below, illustrate the differences in pS129-αSyn accumulations between FR ΔEC and FR EC challenged TG mice. b) Quantitative analysis of pS129-αSyn+ accumulations in the SNpc accounting for two different forms: accumulations cell bodies based on morphological attributes (Cell body) and other forms of accumulations (Other). The majority of accumulations could be attributed to the EC challenge. The combined challenge (FD EC) appears to have a slightly greater impact on pS129-αSyn accumulation. The representative images below illustrate the different observed forms of pS129-αSyn+ accumulations: see details in main text. Stats: Mann-Whitney U test, not corrected for FDR FD, fibre-deprived; FR, fibre-rich (normal chow); PBS, phosphate buffered saline; ΔEC, curli-KO E.coli; EC, wild-type curli expressing E.coli

In the dorsal striatum the FR EC challenged TG mice had the highest levels of pS129-αSyn positive accumulations, with even significant difference to the FR ΔEC (p = 0.018) group (**Fig. 7a**, top panel), as seen in the representative microscopy images (**Fig. 7a**, lower panels). In the SNpc, we saw that the impact of the FD challenge greater than seen in the dorsal striatum (**Fig. 7b**). Both pS129-αSyn positive cell body counts and all “other” pS129-αSyn positive accumulations were especially increased in FD EC challenged mice (“Cell Body”, vs FR ΔEC: p = 0.127; “Other”, vs FD PBS: p = 0.109, vs FR ΔEC: p = 0.019, vs FR EC: p = 0.072, vs FD ΔEC: p = 0.13, **Fig. 7b**, top panel). Qualitatively, the pS129-αSyn positive accumulations that we observed in cell bodies were usually either of a diffused when in the cytoplasm and more compact in the nuclei (**Fig. 7b**, bottom panel, white arrowhead). However, in FD EC challenged mice we also found dense pS129-αSyn positive accumulations in cell bodies (**Fig. 7b**, bottom panel, blue arrowheads). All other pS129-αSyn positive accumulations, not limited to cell bodies, appeared generally more intensively immunopositive. We observed three different forms of accumulations: bead like varicosities (**Fig. 7b**, bottom panel, white arrow), similar to what has been observed in other *in vivo*(Lauwers et al. 2003) and *in vitro*(Kouroupi et al. 2017) models, and human post-mortem brains(Del Tredici et al. 2002), spheroid shaped accumulations (**Fig. 7b**, bottom panel, open arrow) and, even though rarely and only in FD EC challenged mice, corkscrew-like spheroid accumulations (Fig. 9b, bottom panel, blue arrow). In summary, the EC challenge drove pS129-αSyn positive accumulations and were exacerbated in FD challenged TG mice.

### Combined bacterial curli protein and dietary fibre deprivation challenges drive neurodegeneration in Thy1-Syn14 mice

The loss of neurons in the SNpc and their projections to the dorsal striatum is amongst the main pathological hallmarks of PD(Poewe et al. 2017). To detect neurodegeneration in our TG mice, we stained against tyrosine hydroxylase (TH), an enzyme involved in dopamine synthesis and a marker for dopaminergic neurons and their projections, in both the SNpc and dorsal striatum. Additionally, in the dorsal striatum, we stained for the dopamine transporter (DAT), a marker for dopamine-cycling synapses.

In the dorsal striatum, the combination of FD and EC challenges in TG mice resulted in a significantly reduced area occupied by TH positive projections when compared to the FR ΔEC (p = 0.029) and FR EC (p = 0.04) groups (**Fig. 8a**, top panel). For DAT we observed almost the exact same pattern (**Fig. 8b**, top panel). The FD EC challenged TG mice showed significantly reduced levels in DAT area occupied when compared to FD ΔEC (p = 0.05) and strong trends compared to the FD PBS and FR EC groups (**Fig. 8b**, top panel).

**Fig. 8.**
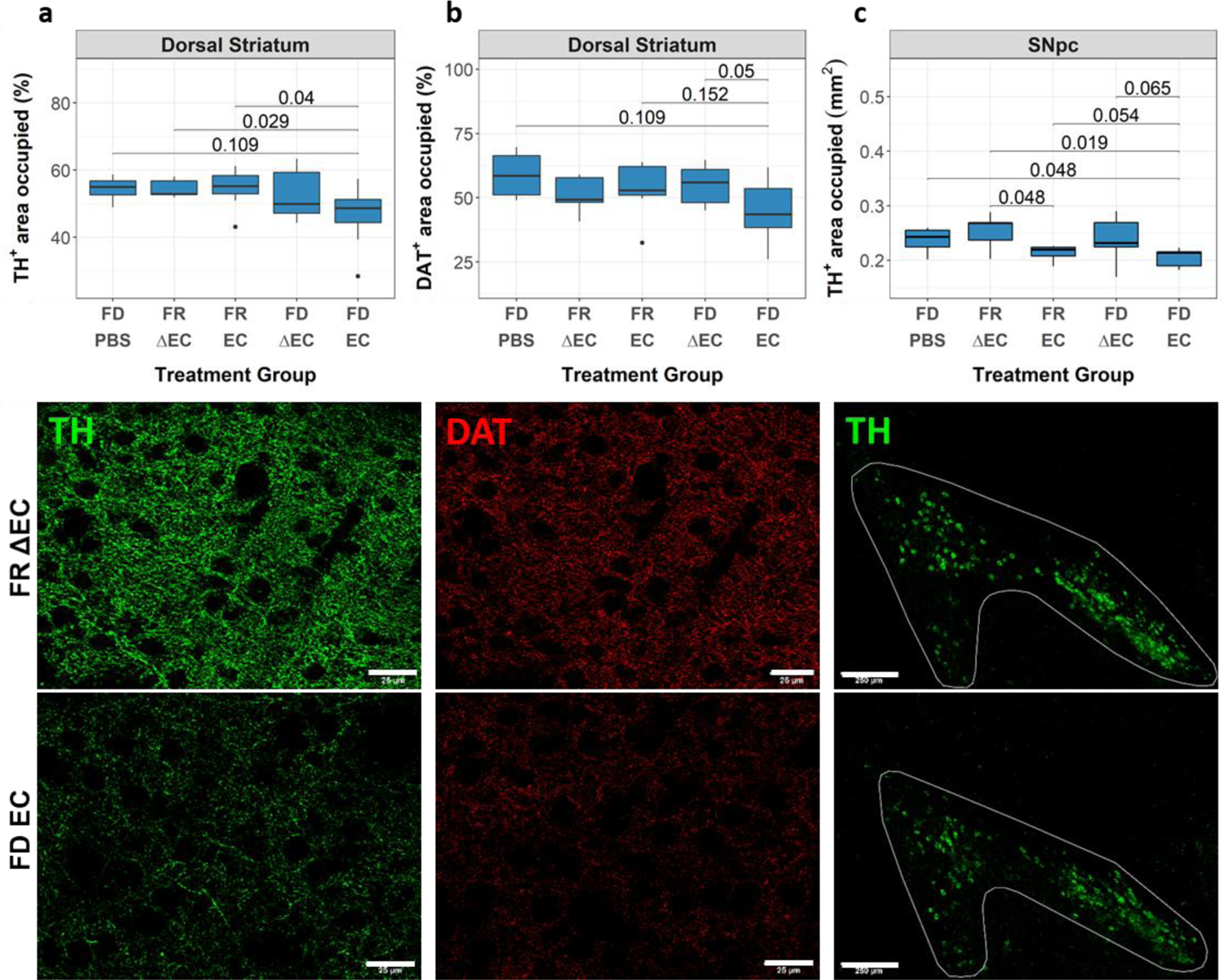
Curli-driven neurodegeneration is exacerbated by fibre-deprivation in the nigrostriatal pathway of Thy1-Syn14 mice. a-c) Boxplots and representative images from two different transgenic treatment groups (top: FR ΔEC; bottom: FD EC) from the dorsal striatum and the substantia nigra pars compacta. Boxplots exhibit the median differences of area occupied by specific neuronal or synaptic markers used to investigate neurodegeneration. a) Quantification and representative high magnification (40X, scale bar: 25µm) images of the percent area occupied by tyrosine hydroxylase-positive (TH+) fibres in the dorsal striatum. b) Quantification and representative high magnification (40X, scale bar: 25µm) images of the percent area occupied by the synaptic dopamine transporter (DAT) marker in the dorsal striatum. c) Quantification and representative images (10X; scale bar: 250µm) of the summed area occupied in square millimeters (mm2) by TH+ dopaminergic neurons in the Substantia Nigra pars compacta. Stats: Mann-Whitney U test, not corrected for FDR SNpc, substantia nigra pars compacta; FD, fibre-deprived (diet); FR, fibre-rich (diet) (normal chow); PBS, phosphate buffered saline; ΔEC, curli-KO E.coli; EC, wild-type curli expressing E.coli

Tyrosine hydroxylase positive fibres in the dorsal striatum are the projections from dopaminergic neurons located in the SNpc. Quantitation of the area occupied by TH positive neurons (**Fig. 8c**, top panel) showed that the exposure to curli caused significant neuronal loss (EC-PBS: FDR = 0.041; EC-ΔEC: FDR = 1.68E-4). Fibre deprived challenged mice, showed an exacerbated neuronal phenotype. The difference between the FR EC and FD EC groups does show a strong (p = 0.054) trend. Hence, the FD challenge potentially leads to an increased susceptibility to a curli-driven neurodegenerative process.

### Summary: The combination of fibre deprivation and bacterial curli exacerbates PD-like pathologies in Thy1-Syn14 mice

To obtain a bird’s-eye view, we generated a radar plot summarising the collection of our findings. We simplified the output by classifying the results of the different treatment groups (FD PBS, FR ΔEC, FR EC, FD ΔEC, FD EC) from lowest (centre of plot, **Fig. 9a**) to highest (plot outline, **Fig. 9a**). We further split our findings over two categories (brain and gut) and 8 sub-categories (brain: Motor impairment, Neurodegeneration and αSyn accumulation (CNS); gut: alpha diversity decrease, mucus foraging genera, lower gut barrier integrity, mucus erosion and αSyn accumulation (ENS); **Fig. 9a**).

**Fig. 9.**
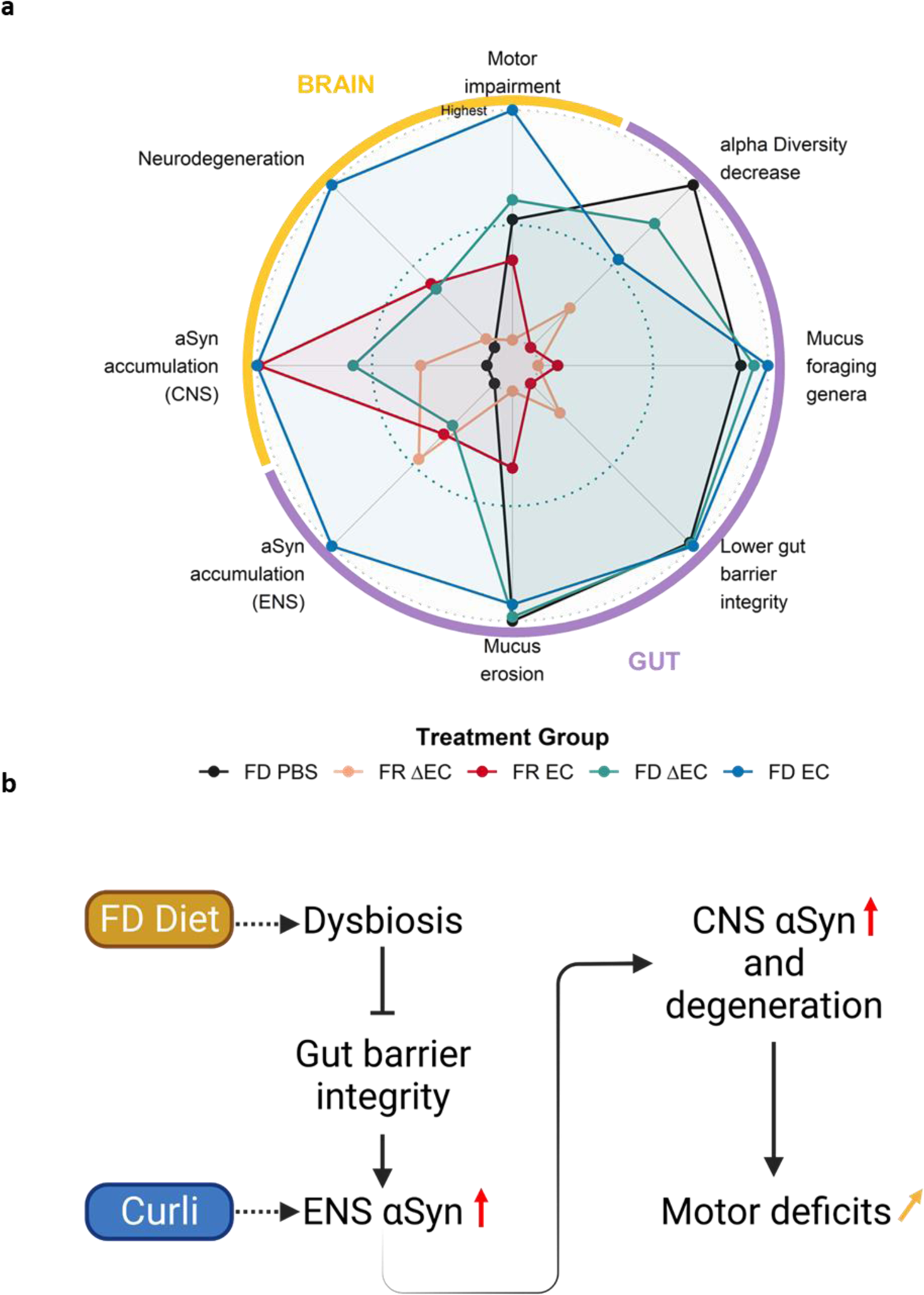
A multi-challenged driven sequence of events for PD progression. a) Radarplot of the total output for each challenge group in Thy1-Syn14 mice. The center of the plot defines the treatment with the lowest and the outline the one that showed the highest effect on Thy1-Syn14 mice. Overall, the combination of FD and EC has the greatest impact on αSyn overexpressing mice. View main text for more details. b) Scheme of a potential sequence of events based on the results in this study. FD challenge leads to changes in the microbiome (dysbiosis), which in turn affects the gut barrier integrity and consequently facilitates the interaction of curli with the submucosa and the plexuses. As a results, αSyn abnormally accumulates. This is further seen in the brain where we observed αSyn accumulation, accompanied by neurodegeneration in the nigrostriatal pathway. these changes impact the locomotion and exacerbate the already strong motor deficits in Thy1-Syn14 mice. CNS, central nervous system; ENS, enteric nervous system; FD, fibre-deprived; FR, fibre-rich (normal chow); PBS, phosphate buffered saline; ΔEC, curli-KO E.coli; EC, wild-type curli expressing E.coli

Under overexpressing αSyn condition, we saw that the diet challenge impacted almost all aspects of the gut and curli drove αSyn accumulation (**Fig. 10**). However, the combination of all challenges (TG FD EC) had the greatest effect, except for alpha diversity decrease, on all PD relevant pathologies.

Based on our results, we propose a sequence of events (**Fig. 9b**) where dietary fibre deprivation led to changes in microbial populations and enteric physiology. These changes cumulated in the increase of gut penetrability. The exposure to the bacterial protein curli caused increased αSyn accumulations in the ENS and in the CNS accompanied by neurodegeneration in the nigrostriatal pathway. These changes finally resulted in the further exacerbation of already strong motor deficits in these TG mice.

## Discussion

Most Parkinson’s disease cases have a complex multi-factorial risk profile. There is only little known on how these varied factors come together to affect or exacerbate PD progression. In our study we investigated how genetically predisposed aged mice were affected by a fibre deprived diet and exposure to curli producing *E. coli*, individually or combined and uncovered a sequence of events that systematically led to the exacerbation of PD pathologies.

There is a great body of evidence on the impact of diet on microbial gut health and how it can influence the course of a disease. *In vivo* studies where rodents were fed a fibre deprived diet saw rapid shifts in microbial gut populations(Schroeder et al. 2018; Desai et al. 2016; Neumann et al. 2021; Riva et al. 2019). Accordingly our data showed decreased diversity, increased *Firmicutes*/*Bacteriodetes* ratios and altered abundance of many taxa. The lack of dietary fibre was shown to make specialized taxa switch to host glycans, which resulted in increased mucus erosion and increased susceptibility to pathogens(Desai et al. 2016; Martens, Chiang, and Gordon 2008). Longer absence of dietary fibre is thought to trigger a compensatory mechanism, increasing mucin production, and re-establishing the inner mucus thickness(Schroeder et al. 2018). Based on our results, however, the outer mucus layer does not recover. The thin outer mucus layer associated with reduced bacterial diversity and therefore it can be assumed that there are changes in host-microbe interactions. Recent studies showed the impact of microbial metabolite changes on gut barrier integrity. The metabolite butyrate for instance is essential in regulating energy metabolism, proliferation, and differentiation of gut epithelial cells, has anti-inflammatory properties, stimulates mucin production, and most importantly is involved in gut barrier protection by stimulating expression of ZO-1, ZO-2, cingulin and occludin(Plöger et al. 2012; Rivière et al. 2016). While we did not assess butyrate levels, we did observe reduced abundances of the butyrate producing genera *Lachnospiraceaea NK4A136* and *Roseburia.* Additionally, *Lactobacillus*, which was also reduced in our fibre deprived fed mice, has been proposed to stimulate butyrate production of such bacteria(Lin et al. 2020). There is still conflicting evidence on the effect of short-chain fatty acids, in particular butyrate, in PD. Most animal studies however report beneficial effects(Paiva et al. 2017; St. Laurent, O’Brien, and Ahmad 2013; Sharma, Taliyan, and Singh 2015). Taken together, even though we do not assess for gut barrier integrity, our observations of reduced outer mucus thickness, higher levels of particular mucus foraging taxa and reduced gut health relevant taxa under fibre deprived conditions did let us conclude that these mice were more susceptible to potential pathogenic factors such as curli.

Curli is a bacterial protein, mainly produced by *Enterobacteriaceae*. This bacterial family has been reported to be increased in PD patients and is associated with disease severity(Barichella et al. 2019; Li et al. 2017). The curli protein has amyloidogenic properties and has been shown to act as a seed for αSyn aggregation *in vitro*(Sampson et al. 2020). In physiological conditions, either gavaged(S. G. Chen et al. 2016) or supplemented in a human faecal microbiota transplant(Sampson et al. 2020), curli presence led to different PD pathologies. Chen and colleagues observed increased αSyn accumulation in the gut of exposed Fischer 344 rats(S. G. Chen et al. 2016). Interestingly, the deposits observed in the gut were soluble while they did find proteinase K resistant αSyn, as one would find in Lewy bodies, in the brain(S. G. Chen et al. 2016). Analogously, we found increased levels of pS129-αSyn in the gut, more specifically in the myenteric plexus. The other study by Sampson and colleagues did not investigate αSyn in the ENS, but they did show increased pS129-αSyn positive levels in the brain in transgenic mice exposed to curli producing *E. coli*. Neither these studies nor we elucidate on the spreading mechanism from the ENS to the CNS via the vagus nerve. The spreading hypothesis was postulated by Braak and colleagues after they discovered that in some patients, αSyn deposits preceded CNS pathologies. The spreading process is described as “prion-like” since pathological forms of αSyn act as seed for yet uncorrupted αSyn (Jucker and Walker 2018; Mezias et al. 2020), similar to what was proposed for curli (Chapman et al. 2002; Barnhart and Chapman 2006; Sampson et al. 2020). Based on post-mortem observations, the Braak hypothesis posits that αSyn first accumulates in the lower regions of the brainstem namely the dorsal motor nucleus of the vagus. Subsequently, αSyn deposits gradually move upwards in a “prion-like” manner, resulting in different PD symptoms from early non-motor to the typical motor symptoms as the disease progresses. Direct evidence for the “prion-like” spreading comes from both *in vitro* and *in vivo* studies(Rey et al. 2016; Vasili, Dominguez-Meijide, and Outeiro 2019). Animal models which were injected directly into the brain with different forms of αSyn developed a variety of PD pathologies including spreading of pathological endogenous αSyn(Rey et al. 2016; Luk et al. 2012; Garcia et al. 2022). To investigate whether the propagation via the vagus is a possible route, a team from Johns Hopkins University injected preformed αSyn fibrils into the muscle layers of the GI tract of non-transgenic mice(Kim et al. 2019). They observed a progressive retrograde propagation originating in the dorsal motor nucleus of the vagus. After three months, the SNpc showed pS129-αSyn positive accumulations, followed by significant degeneration at month seven(Kim et al. 2019). After truncal vagotomy, both αSyn deposits and neurodegeneration were absent(Kim et al. 2019). This is in accordance with epidemiological meta-analyses suggesting that truncal vagotomy reduces the risk to develop PD.

Taken together, this study supports the idea of the gut-brain-axis in PD and thus Braak’s spreading hypothesis. To our knowledge we are the first to propose a combinatorial mechanism of interdependent exogenous and endogenous factors contributing to the progression PD. Additionally, we cannot exclude that these events are also involved in the onset of PD. We further underlined the importance of a balanced healthy diet and its implications in disease progression. Hence, our results propose a translational PD-relevant sequence of events putting forth the idea for lifestyle adaptations to prevent or mitigate disease progression.

## Material and Methods

### Animals and experimental design

#### Ethical Approval

All animal experimentations were approved by the Animal Experimentation Ethics Committee of the University of Luxembourg and the appropriate Luxembourg governmental agencies (Ministry of Health and Ministry of Agriculture) and registered under LUPA 2020/25. Additionally, all experiments were planned and executed following the 3R guidelines (https://www.nc3rs.org.uk/the-3rs) and the European Union directive 2010/63/EU.

#### Mice

We used the transgenic line B6.D2-Tg(Thy1-SNCA)14Pjk(Philipp J. Kahle et al. 2000; P. J. Kahle et al. 2001), which we will refer to as Thy1-Syn14 or TG from here on forth. This line overexpresses wild-type human αSyn under the transcriptional regulation of the neuron specific Thy1 promoter. As control animals we used the wild-type (WT) littermates. All mice used were male. For the characterization of the line we used four different cohorts. For the experimental challenge cohort, we used 72 male animals, 36 TG and 36 WT littermates. They were singly-caged to avoid coprophagy, had access *ad libitum* to food and water and were exposed to a regular 12h-day-night cycle. Animals were monitored twice a week. According to our welfare guidelines, we set a humane endpoint based on different physical parameters, e.g. weight loss/gain, body temperature and coat condition. At the end of the in-life phase, we anesthetized the mice with a mix of 150mg/kg ketamine + 1mg/kg medetomidine, collected blood from the right atrium and subsequently transcardially flush-perfused with 1X PBS.

During the in-life phase of the study, 10 mice were either found dead in their home cage or reached a humane endpoint (**Supplementary Fig. 1a**). This measure agrees with the animal welfare guidelines.

#### Experimental Design

For the challenge study, mice were randomly assigned to 10 different treatment and respective control groups (**Supplementary Fig. 1a**) and treated for a total of 9 weeks; 1 week dietary priming of the colon and additional 8 weeks combined diet and bacterial challenges (**Supplementary Fig. 1b**). We provided fresh diet and gavaged the animals with the respective bacteria or sham solution weekly. Body weight and overall health was checked twice a week. We collected stool samples for microbiome analysis and monitored basis gross motor functions by assessing hind limb clasping and grip strength weekly (**Supplementary Fig. 1b**). After euthanasia we collected brains and colons for molecular biology and histology.

### Bacterial solution preparation and gavage

The *E. coli* strains used for treatment were the C600 (EC) and its isogenic curli-operon knock-out (ΔEC) strains(Chapman et al. 2002; S. G. Chen et al. 2016). Both strains were a kind gift from Matthew Chapman, University of Michigan. Expression/absence of curli operon was tested via PCR using the following primer pairs: *csgA*_F-5’-GCGTGACACAACGTTAATTTCCA-3’, *csgA*_R-5’-CATATTCTTCTCCCGAAAAAAAACAG-3’; *csgB*_F-5’-CCATCGGATTGATTTAAAAGTCGAAT-3’, *csgB*_R-5’-AATTTCTTAAATGTACGACCAGGTCC-3’. Additionally, curli protein expression was confirmed by Congo red staining (not shown). Both strains were grown in Lennox broth under aerobic (5% CO_2_) conditions at 37°C agitating at 300rpm. Bacteria were resuspended in sterile PBS for oral administration at 10^10^CFUs/mL.

They were gavaged 100µL of bacterial solution, or PBS for the gavage control groups, at a total bacterial load of 10^9^CFUs. Reusable stainless steel 20G feeding needle (Fine Science Tools, 18060-20) were used. Prior and in-between gavages, the feeding needle was washed with filtered 70% ethanol and rinsed with sterile PBS. One group of feeding needles per treatment were used to avoid cross-contamination. Importantly, we did not pre-treat our mice with an antibiotic mix because 1) antibiotics have been shown to prevent αSyn aggregation and to be neuroprotective(Yadav et al., n.d.), and 2) we were interested in the impact of the fibre deprivation on a native microbiome.

### Tissue collection and preparation

Prior to the transcardial perfusion, we collected up to 400µL of venous blood from the right atrium in EDTA K3 coated collection tubes (41.1504.005, Sarstedt), for blood-endotoxin measurements. The tubes were gently inverted and then kept on ice. The plasma was collected after centrifugation at 2000 x g for 10 mins, transferred to RNase-free tubes and stored at −80°C.

After perfusion, the brain was placed on ice and split along the longitudinal fissure into two hemibrains. For molecular biology analyses, one hemibrain was dissected into different regions of interest (striatum and ventral midbrain). The dissected regions were then put on dry ice and further stored at −80°C. The other hemibrains were fixed for immunohistochemistry in 4% PBS-buffered paraformaldehyde (PFA) for 48h at 4°C and then stored in PBS-azide (0.02%) at 4°C. Subsequently, they were cut to generate 50µm thick free-floating sections using the Leica vibratome VT1000 (Wetzlar, Germany) and stored the sections in a 1% (w/v) PVPP + 1:1 (v/v) PBS/ethylene glycol anti-freeze mix at −20°C until staining.

Colon samples for mucus measurements and histopathology were fixed in methacarn (60% absolute methanol: 30% chloroform: 10% glacial acetic acid) solution for 2-4 hours, then transferred to 90% ethanol and kept at 4°C. Next, whole colon samples were first transversally cut by hand with a microtome blade into 4-5mm long sections. Those pre-cut sections were then put into a histology cassette while respecting the proximal to distal order. They were held in place in an ethanol soaked perforated sponge. After 24h post-fixation in 10% formalin, the samples were processed in a vacuum infiltration processor. Finally, all samples were embedded in paraffin and cut at 3µm on a microtome. If not processed immediately, the slides were stored at 4°C before the stainings.

### RNA extraction and RT-qPCR

RNA from dissected colon and different brain regions was extracted using the Qiagen RNeasy Plus Universal Mini Kit (Qiagen, 73404). Briefly, 900µL QIAzol lysis buffer (Qiagen, 79306) and three cold 5mm steel balls were added to each sample (previously stored at −80°C). In ice cooled racks, samples were homogenized at 20Hz for 2mins using the Retsch Mixer Mill MM400. Homogenates were transferred to new RNase clean 2mL tubes and left to rest for 5mins at room temperature (RT). 100μL gDNA eliminator solution was added and the tubes were shaken vigorously for 15 seconds. Then 180μL of chloroform was added and another strong shake was applied for 15 seconds. Homogenates were left to incubate for 3mins at RT. Samples were then centrifuged at 12000 x *g* for 15 mins at 4°C. Five hundred μL of the upper aqueous phase was collected and transferred to new 2mL RNase free tubes. Five hundred μL of ethanol was added to the supernatant and mixed by inverting the tubes back and forth. RNeasy mini spin columns were then loaded with 500μL of the mix, centrifuged at 8000 x *g* for 30s at RT, followed by discarding the flow-through from the collection tube. This step was repeated once more. The spin columns were then washed with two different buffers in three steps: one time with 700μL of RWT buffer and twice with 500μL of RPE buffer. At each washing step the columns were centrifuged at 8000 x *g* for 30s at RT and the flow-through was discarded. Columns are then transferred to new collection tubes and spun at maximum speed for 1min. Finally, columns were transferred to an RNase free 1.5mL Eppendorf tube. Fifty μL of RNase-free water was added to the columns to elute total RNA. RNA purity and quantity were checked by spectrophotometry using the NanoDrop™ 2000 (ThermoFisher Scientific) and the Agilent 2100 Bioanalyzer, respectively. Finally, RNA samples were stored at −80°C.

The model used in this study, Thy1-Syn14, carries a transgene for wild-type human αSyn (*SNCA*). To determine the levels of transcript expression in comparison to endogenous murine αSyn (*Snca*), quantitative RT-PCR was performed on a separate untreated age-matched male cohort (N=15), using the following primer pairs: Snca F 5’-GAT-CCT-GGC-AGT-GAG-GCT-TA-3’, R 5’-CT-TCA-GGC-TCA-TAG-TCT-TGG-3’, SNCA F 5’-AAG-AGG-GTG-TTC-TCT-ATG-TAG-GC-3’, R 5’-GCT-CCT-CCA-ACA-TTT-GTC-ACT-T-3’ and reference gene *Gapdh* F 5’-TGC-GAC-TTC-AAC-AGC-AAC-TC-3’, R 5’-CTT-GCT-CAG-TGT-CCT-TGC-TG-3’. For the reverse transcription of RNA to cDNA we used the SuperScript™ III RT reverse transcriptase from Invitroge. Briefly, 1μL of oligo (dT) 20 (50 μM) and 1μL of 10mM dNTP mix was added to 1μg of total RNA. If needed, nuclease free water was added to obtain the final reaction volume of 13µL. The mixture was briefly centrifuged for 2-3s, heated at 65°C for 5 minutes, and again chilled on ice for at least 1 minute. Another mixture of 4μL 5× first strand buffer, 1μL RNaseOUT (RNase inhibitor), 1μL of 0.1M DTT and 1μL of Superscript reverse transcriptase (200 U/μl) was added. The final mixture was briefly centrifuged and incubated at 50°C for 1h followed by 15mins at 70°C for enzyme deactivation. 80μL of RNase free water was added to the reaction mixture. The obtained cDNA was then placed on ice for immediate use or stored at −20°C for future use.

The qPCR reaction mix contained 2µL of cDNA, 10µM forward and reverse primers, 1X iQ^TM^ SYBR® Green Supermix (Bio-Rad) and PCR grade water up to a volume of 20µL. Each qPCR reaction was run in duplicates on a LightCycler ® 480 II (Roche). The thermo cycling profile included an initial denaturation of 3 minutes at 95°C, followed by 40 Cycles at 95°C for 30 seconds, 62°C (annealing) for 30 seconds and 72°C (elongation) for 30 seconds, with fluorescent data collection during the annealing step. Data acquisition was performed by LightCycler® 480 Software (version 1.5.0.39).

### Microbial DNA extraction and 16S rRNA Amplicon Sequencing

For microbial DNA extraction from single faecel pellets we used an adapted version of the IHMS protocol H(Dore et al. 2015). Faecal samples were preserved in 200µL of a glycerol (20%) + PBS solution and stored at −80°C. Prior to the extraction, we slightly thawed the samples and added 250µL guanidine thiocyanate and 40µL N-lauryl sarcosine (10%). The samples were then left at RT to fully thaw. We then added 500µL N-lauryl sarcosine (5%) before the faecal pellet was scattered and vortexed to homogeneity. Samples were then shortly spun down and incubated at 70°C for 1h. Seven hundred fifty µL pasteurized zirconium beads were added to the tubes, then put in pre-cooled racks and horizontally shaken for 7.5mins at 25Hz in a Retsch mixer mill MM400. Fifteen mg polyvinylpyrrolidone (PVPP) was added and vortexed until dissolved. Then the samples were centrifuged at 20814 x g for 3mins. The supernatants were transferred to new 2mL tubes and kept on ice. We then washed the pellet with 500µLTENP (Tris, EDTA, NaCl and PVPP) and centrifuged at 20814 x g for 3mins. This step was repeated three times in total and each supernatant was added to the previously new 2mL tube. To minimize carryover, the tubes were centrifuged again at 20814 x g for 5mins and the supernatant was split equally in two new 2mL tubes. We then added 1mL isopropanol (Merck) to each tube and mixed them by inverting the tubes. After a 10min incubation at RT, the samples were centrifuged at 20814 x g for 15mins. The supernatant was discarded and the remaining pellet air dried under the fume hood for 10mins. The pellet was then resuspended in 450μL phosphate buffer and 50μL potassium acetate by pipetting up and down, before the duplicates were pooled and incubated on ice for 90mins. Then the sample was centrifuged (20,814 x g) at 4 °C for 35mins, the supernatant transferred into a new tube. Next, 2μL of RNase (10 mg/ml) were added. Then, the tube was vortexed, briefly centrifuged and finally incubated at 37 °C for 30mins. We then added 50μL of sodium acetate, 1mL of ice cold 100% ethanol (Merck) and mixed the tube by inverting several times. The sample was again incubated at RT for 5mins and centrifuged at 20814 x g for 7.5 min. The supernatant was discarded and the newly formed pellet was subsequently washed three times in total with 70% ethanol (Merck) and centrifuged at 20814 x g for 5mins. The supernatant was discarded each time. Finally, the clean pellet was dried at 37 °C for 15 min, then resuspended in 100μl TE Buffer and homogenized by pipetting. After incubation at 4°C over night, DNA quality and quantity were checked by Nanodrop ™ 2000/2000c and Qubit 2.0 fluorometer (Thermo Fischer Scientific). Samples were stored at −80°C until sequencing.

Five ng of isolated gDNA were used for PCR amplification using primer (515F (GTGBCAGCMGCCGCGGTAA) and 805R (GACTACHVGGGTATCTAATCC)) specific to V4 region of 16S rRNA gene. For the first round of PCR, samples were amplified for 15 cycles to avoid over-amplification. Additional 6 PCR amplification cycles were performed in the second round to introduce sample specific barcode information. All samples were pooled in equimolar concentration for sequencing. Sample preparation and sequencing were performed at LCSB Sequencing platform using v3 2×300 nucleotide paired end sequencing kit for MiSeq.

### 16S rRNA gene amplicon sequence analysis

#### Sequence analysis

Amplicon Sequence Variants (ASVs) were inferred from 16S rRNA gene amplicon reads using the dada2 package(Callahan et al. 2016) following the paired-end big data workflow (https://benjjneb.github.io/dada2/bigdata_paired.html, accessed: September, 2020), with the following parameters: truncLen = 280 for forward, 250 for reverse reads, maxEE = 3, truncQ = 7, and trimLeft = 23 for forward, 21 for reverse reads. The reference used for taxonomic assignment was version 138 of the SILVA database (https://www.arb-silva.de)(Quast et al. 2013).

#### Microbial diversity and related statistics

Microbiome count data was managed using the phyloseq R package(McMurdie and Holmes 2013) this package was also used to calculate the Shannon index for alpha diversity and non-metric multidimensional scaling (NMDS) ordination for beta diversity. Statistical significance of alpha diversity differences was evaluated with the Kruskal-Wallis test (overall comparison between all groups) and the Wilcoxon Rank Sum Test with false discovery rate correction for multiple comparisons (pairwise contrasts). For beta diversity comparisons, we used the adonis PERMANOVA test from the R package vegan (Oksanen et al. 2020). All diversity comparisons were performed using ASV count data rarefied to the lowest number of sequences in a sample. Taxon-specific plots (genus and family level) were made using relative abundances (% of taxa out of total).

### Endotoxin plasma level measurement by ELISA

Plasma samples were diluted 1:10 in 1X PBS. To measure the endotoxin plasma we used the EndoLISA® kit from BioVendor. We followed the supplier’s protocol. Briefly, a serial dilution for the standard for the non-linear regression model was prepared. In duplicates, 100µL of well mixed standard and samples were applied on the supplied 96-well plate. Then, 20µL of 6X binding buffer was added to each well and the plate was sealed with a cover foil. The plate was incubated at 37°C for 90 mins on a shaker at 450rpm. The reaction buffer was removed by quickly inverting the plate. The excess buffer was removed as best as possible by tapping the plate gently on a paper towel. The wells were then washed twice with 150µL of wash buffer. Again, the plate was inverted quickly and the excess buffer was removed by tapping the plate gently on a paper towel. Finally, we added 120µL of Detection buffer and start measurements in a 37°C pre-heated plate reader. We measured from T=0min to T=90mins at 15mins intervals.

### Alcian blue staining and outer mucus thickness measurements

The Alcian blue stainings were performed at the National Center of Pathology (NCP) of the Laboratoire National de Santé in Dudelange (Luxembourg). The sections were stained for Alcian blue (Artisan Link Pro Special Staining System, Dako, Glostrup, Denmark) according to manufacturer’s instructions.

We took 5-10 images per section at 20X. This resulted in up to 24 images per animal. The criteria for the correct images were that the sections were cut at the correct plane level. This was determined by the orientation and definition of the crypts, which had to fully visible pointing towards the colonic lumen. We measured only outer mucus areas which could clearly be distinguished from the inner mucus layer and the colonic content. After the images were scaled in image using the imprinted scale bar as reference, an average of 6 measure points, spanning the outer mucus layer, per image were taken using Image J.

### Immunofluorescent staining of colon sections

We followed a standard protocol with minor adjustments(‘Immunofluorescent Staining of Paraffin-Embedded Tissue’ n.d.). Briefly, sections were deparaffinised in xylene 3 x 5mins. Before proceeding to rehydration, we checked that all paraffin was removed. If not, the sections were treated another round with xylene. A three step rehydration step with 100%, 70% and 50% ethanol followed deparaffinization. After washing with dH_2_O, we proceeded with a citrate buffer (0.1M, pH6.0, + 0.1% Tween 20) based antigen retrieval at 80°C for 30-35mins. After letting the sections cool down for 20mins, the slides were again washed with dH_2_O 2 x 5mins. We proceeded as described previously(‘Immunofluorescent Staining of Paraffin-Embedded Tissue’ n.d.) for peroxidase inactivation, permeabilization and blocking. After washing again with dH_2_O, the tissue was circled with a hydrophobic Dako pen (S2002, DAKO) and the primary antibodies were added. They were incubated at room temperature (RT) for 2 hours (hrs) and then transferred to 4°C for overnight incubation in a humidified chamber. The following day, slides were washed briefly with dH_2_O and then washed with 1% BSA + PBS 0.4% Triton X100 2 x 5mins and finally rinsed with dH_2_O. Tissues were circled again with the hydrophobic pen and secondary antibodies were added. Slides were then incubated for 2 hrs at RT in the humidified chamber. Finally, they were washed 3 x 5mins with PBS 0.4% Triton X100 and rinsed with dH_2_O. Excess water was removed by gentle tapping and slides were coverslipped with DAPI Fluoromount-G® (0100-20, SouthernBiotech).

To detect phosphorylated αSyn in the ENS we performed double staining using the following antibodies: polyclonal chicken anti-PGP9.5 (ab72910, Abcam; 1:1000), monoclonal rabbit anti-pS129-αSyn (ab51253, Abcam; 1:500).

### Behaviour

#### Hindlimb Clasping

The method was adapted from Bouet et al.(Guyenet et al. 2010). Brief, animals were taken by the tail near the base and suspended for 10 seconds. If both hindlimbs stayed stretched and did not touch the abdomen for more than 50% of the suspension time we scored it 0. A score of 1 or 2 was given if one respectively both hindlimbs were retracted for more than 50% of the suspension time. If they were retracted and touched the abdomen for the entire suspension time a score of 3 was given. In the most severe cases, the animals twisted around the vertical body axis or even rolled up to a so-called bat position. These cases were given a score of 4. This test was repeated weekly.

#### Grip Strength

The grip strength test(Mao et al. 2016) was performed using Bioseb’s grip strength meter (Vitrolles, France). Animals were gently placed on a grid, allowed to grab onto it with all four paws and then gently pulled off in a continuous backwards motion by their tail. Technical triplicates were taken for each mouse. Values were normalized to the weight of the respective mouse. The test was repeated weekly.

#### Adhesive Removal

The test was adapted from Bouet and colleagues (Bouet et al. 2009). Brief, animals were placed in a round transparent arena for one minute as habituation. A piece of rectangular tape (3×5mm) was placed on each forepaw. The time was taken once the animals touched the bottom of the arena. We measured the time of first touch and first removal. The test was performed in duplicates, and performed at baseline and at the end of the experimental phase.

### Immunofluorescent staining on free-floating brain sections

Immunofluorescent stainings on free-floating sections were performed following a standard protocol(Ashrafi et al. 2017) with minor adaptations. Briefly, sections were washed in PBS + 0.1% Triton X100 (T_X100_) to rinse off the anti-freeze solution. Then, they were treated with a permeabilization/peroxidase inactivation solution (PBS + 1.5% T_X100_ + 3% H_2_O_2_) for 30mins followed by 2×5mins washing. To prevent unspecific antibody binding, the sections were incubated in 5% BSA + 0.02% T_X100_ for 1 hour. After a short washing step, sections were incubated with primary antibody(ies) diluted in antibody solution (PBS + 2% BSA) over night at room temperature (RT) on an orbital shaker. The next day, sections were washed with PBS + 0.1% TX100 to remove all excess first antibody. Sections were then incubated with secondary antibody (+ antibody solution) for 2 hours at RT on an orbital shaker under a light trap. Finally, sections were washed with simple PBS (at least three times for 10mins) and then mounted on Superfrost™ (ThermoFisher Scientific) slides, let to dry for up to 12h, and cover-slipped using the Fluoromount-G® (Invitrogen) mounting solution.

The following antibodies were used: monoclonal rabbit anti-pS129-αSyn (Abcam, ab51253; 1:1000), monoclonal mouse anti-pS129-αSyn (Prothena Biosciences Inc., 11A5; 1:1000), polyclonal chicken anti-tyrosine hydroxylase (Abcam, ab76442; 1:1000), polyclonal rabbit anti-tyrosine hydroxylase (Merck (Sigma-Aldrich), AB152; 1:1000), polyclonal rat anti-dopamine transporter (MAB369, Merck (Sigma-Aldrich); 1:1000).

### Quantitative neuropathology

We performed the imaging our sections using a Zeiss AxioImager Z1 upright microscope, equipped with a PRIOR motorized slide stage and coupled a “Colibri” LED system to generate fluorescence light of defined wavelengths, and a Zeiss Mrm3digital camera for image capture. The complete imaging system was controlled by Zeiss’ Blue Vision software. All histological analyses were performed blinded.

The quantification of TH-positive fibres and DAT-positive synaptic terminals was done as described before in Garcia *et al*., 2022(Garcia et al. 2022). Briefly, two doubly labelled (rabbit anti-TH and rat anti-DAT) sections were used. A total of 6 (3/section) 40x (223.8 × 167.7 μm^2^) pictures of the dorsal striatum were acquired using the optical sectioning system Apotome.2 (Zeiss). The percent area occupied of TH and DAT by intensity thresholding was determined using Image J software and averaged for each mouse.

The quantification of TH-positive neurons in the SNpc has been described and the obtained results have been correlated with stereological cell counts (see supplementary information in Ashrafi et al., 2017(Ashrafi et al. 2017)). Briefly, to estimate TH-positive neurons in the SNpc, anatomically distinguishable levels were identified and applied to 7-12 fifty-micron sections/mouse. Then, 2×2 tiled pictures/section were taken at 10X objective and converted into single Tiff files for image analysis. Next, the region-of-interest (ROI) of the area occupied only by TH-positive neurons was outlined. After thresholding, the ROI occupied (in pixels) by TH-positive neurons was measured. For each anatomical levels of the SN, up to 2 sections/level were measured. Single and/or averaged values/level were finally summed up to one single representative value, the “cumulative SN surface” and converted to mm^2^.

To quantify pS129-αSyn in the dorsal striatum (Double label: area reference marker polyclonal rabbit anti-TH; pS129-αSyn marker monoclonal mouse 11A5), 20X tile images were converted to 8-bit, the ROI was determined and the threshold was automatically set to “MaxEntropy”. Next, the images were appropriately scaled from pixel to m. This allowed us to adjust our settings to exclude all particles surpassing the size of a synapse. These settings in “Analyze Particles” were *Size (µm2): 30.00-Infinity; Circularity: 0.25-1*. Each particle created an enumerated ROI and was added to the ROI Manager. All images were then manually curated to delete falsely selected or to add missed particles. Subsequently, the full picture frame and all ROIs to be excluded from quantification were selected and combined by “XOR”. Finally, the percentage area occupied and intensity after thresholding for pS129-αSyn positive synaptic areas were measured.

To estimate pS129-αSyn positive accumulations in the SNpc (monoclonal rabbit anti-pS129-αSyn), the same ROIs as chosen for TH quantification were used. In Image J, a virtual grid with a square area of 2500µm^2^ was overlaid. Pictures were then manually analysed for pS129-αSyn positive accumulations. We counted separately, based on morphology, pS129-αSyn positive cell bodies (1 count = 1 cell body) and other pS129-αSyn positive particles (number of particles per square). Finally, counts were normalized per square (area of ROI/area of square) and summed up for all 4 zones (see TH quantification). Phosphorylated-αSyn intensities in the SNpc were not measured.

### Statistics

For the characterisation of the Thy1-Syn14 to test for differences in gene expression levels, protein levels and motor behaviour performance we used the Kruskal-Wallis test and corrected for FDR. If needed we adjusted for multiple comparison using the Mann-Whitney U test and corrected for FDR. For the statistics of the 16S amplicon rRNA sequencing data see above. For all other measurements of the challenged mouse cohort, we performed the non-parametric Kruskal-Wallis test and post-hoc Mann-Whitney U test to adjust for multiple comparison. We additionally corrected for FDR. For all other experiments we used the Mann-Whitney U test to adjust for multiple comparison. If corrected or not for FDR is specified in the figure legends.

## Data Availability

All original datasets are available upon reasonable request to the corresponding author.

## Acknowledgments

Kristopher J. Schmit was recipient of a pre-doctoral fellowship (FNR AFR 12515776) from the Luxembourg National Research Fond. Alessia Sciortino is part of PARK-QC DTU funded by the Luxembourg National Research Fund (PRIDE17/12244779/PARK-QC). Michel Mittelbronn thanks the Luxembourg National Research Fond for support (FNR PEARL P16/BM/11192868). The authors thank Wagner Zago (Prothena Biosciences) for providing the 11A5 antibody and Matt Chapman (University of Michigan) for providing the two *E. coli* strains used in this study. The authors would like to thank the Animal Facility of the University of Luxembourg for their support throughout the animal experiments.

## Abbreviations

PD: Parkinson’s disease

αSyn: alpha-synuclein

pS129-αSyn: phosphorylated S129 alpha-synuclein

TG: Thy1-Syn14 transgenic mice

WT: wild-type littermates

FR: fibre-rich or normal chow

FD: fibre-deprived diet

EC: wild-type *E. coli* expressing curli protein

ΔEC: curli-operon KO *E. coli* strain

## Additional information

### Ethics approval

Animal studies performed at the Luxembourg Centre for Systems Biomedicine were approved by the institutional Animal Experimentation Ethics Committee of the University of Luxembourg, and the responsible Luxembourg government authorities (Ministry of Health, Ministry of Agriculture), following EU directive 2010/63/EU.

### Consent for publication

All authors have approved of the contents of this manuscript and provided consent for publication.

### Availability of materials

The 11A5 monoclonal anti α-synuclein antibody can be obtained, under an MTA, from Prothena Biosciences.

### Authors contributions

K.J.S., M.B., E.C.M. and P.W. designed the study. K.J.S, A.S., B.P.R., P.G., M.H.T., J.J.G., I.B.A., C.C., and T.H did the experiments (gavages, behavioural tests, tissue processing, stainings, imaging, DNA, RNA and protein extractions, Western blots). R.H. performed the 16S rRNA amplicon sequencing. V.T.E.A. analysed the 16S rRNA amplicon sequencing data. K.J.S., A.S., V.T.E.A., P.G., I.O., M.M., M.B., and P.W. analyzed and interpreted the data. K.J.S. drafted the paper. All authors read and approved the final manuscript.

## Notes

### Competing Interest Statement

The authors have declared no competing interest.

