## Supplementary Material for "Dietary fibre deprivation and bacterial curli exposure shift gut microbiome and exacerbate Parkinson’s disease-like pathologies in an alpha-synuclein-overexpressing mouse"

***Supplementary Table 1:*** *Comparison to known taxa altered in stool samples of PD patients (adapted from Boertien et al., 2019)*

| | | | | FD PBS | | | | | | FR ΔEC | | | | | | FR EC | | | | | | FD ΔEC | | | | | | FD EC | | | | | |
|---|---|---|---|---|---|---|---|---|---|---|---|---|---|---|---|---|---|---|---|---|---|---|---|---|---|---|---|---|---|---|---|---|---|
| | | | | 2 | | 5 | | 9 | | 2 | | 5 | | 9 | | 2 | | 5 | | 9 | | 2 | | 5 | | 9 | | 2 | | 5 | | 9 | |
| | | | | WT | TG | WT | TG | WT | TG | WT | TG | WT | TG | WT | TG | WT | TG | WT | TG | WT | TG | WT | TG | WT | TG | WT | TG | WT | TG | WT | TG | WT | TG |
| | **Phylum** | **Family** | **Genus** | | | | | | | | | | | | | | | | | | | | | | | | | | | | | | |
| **Increased** | Verrucomicrobia | | | ↑ | ↑ | ↗ | - | ↑ | ↗ | ↓ | ↓ | - | ↓ | - | ↓ | - | ↓ | ↓ | ↓ | ↓ | ↓ | ↑ | ↑ | ↑ | ↑ | ↑ | ↑ | ↓ | ↑ | ↓ | ↑ | ↓ | ↑ |
| | Verrucomicrobia | Akkermansiaceae | | ↗ | ↑ | ↗ | - | ↗ | ↗ | - | ↓ | - | ↓ | - | ↓ | - | ↓ | - | ↓ | - | ↓ | ↗ | ↑ | ↗ | ↗ | ↑ | ↑ | ↘ | ↑ | ↓ | ↑ | ↓ | ↗ |
| | Verrucomicrobia | Akkermansiaceae | Akkermansia | ↑ | ↑ | ↗ | - | ↑ | ↑ | ↓ | ↓ | - | ↓ | - | ↓ | ↗ | ↓ | ↓ | ↓ | ↓ | ↓ | ↗ | ↑ | ↗ | ↑ | ↑ | ↑ | ↓ | ↑ | ↓ | ↑ | ↓ | ↗ |
| | Firmicutes | Lactobacillaceae | | - | - | - | - | - | - | ↘ | - | ↘ | ↗ | - | ↑ | ↗ | ↑ | ↗ | ↑ | ↑ | ↑ | ↓ | - | ↓ | - | ↓ | - | - | - | - | - | - | - |
| | Firmicutes | Lactobacillaceae | Lactobacillus | ↗ | ↘ | ↓ | ↓ | - | - | ↓ | - | ↘ | ↗ | - | ↑ | ↗ | ↑ | ↗ | ↑ | ↑ | ↑ | ↓ | ↘ | ↓ | ↓ | ↓ | - | ↓ | - | ↓ | ↓ | ↓ | - |
| | Actinobacteria | Bifidobacteriaceae | | *na* | | | | | | *na* | | | | | | *na* | | | | | | *na* | | | | | | *na* | | | | | |
| | Actinobacteria | Bifidobacteriaceae | Bifidobacterium | *na* | | | | | | *na* | | | | | | *na* | | | | | | *na* | | | | | | *na* | | | | | |
| | Proteobacteria | Enterobacteriaceae | | *na* | | | | | | *na* | | | | | | *na* | | | | | | *na* | | | | | | *na* | | | | | |
| | Firmicutes | Enterococcaceae | | *na* | | | | | | *na* | | | | | | *na* | | | | | | *na* | | | | | | *na* | | | | | |
| **Decreased** | Firmicutes | | | ↑ | - | ↗ | ↑ | ↑ | ↑ | - | ↓ | ↗ | - | ↘ | - | ↗ | - | ↗ | ↗ | ↗ | ↗ | ↘ | - | ↘ | ↗ | ↑ | ↑ | - | ↘ | - | ↘ | ↑ | ↗ |
| | Firmicutes | Lachnospiraceae | | ↗ | - | - | ↑ | ↑ | - | ↗ | ↘ | ↗ | - | ↘ | ↘ | ↗ | - | ↗ | ↗ | - | ↗ | ↘ | ↘ | ↘ | ↘ | - | ↘ | - | ↘ | ↗ | - | ↓ | - |
| | Firmicutes | Lachnospiraceae | Roseburia | ↗ | - | ↗ | - | ↗ | - | ↗ | ↘ | ↗ | ↘ | ↗ | ↘ | - | - | - | - | ↑ | ↑ | - | - | - | ↑ | ↑ | ↑ | ↓ | ↘ | ↘ | ↘ | ↘ | - |
| | Bacteroidetes | Prevotellaceae | | ↘ | ↘ | ↘ | ↘ | ↘ | ↘ | - | ↑ | - | - | - | - | - | ↗ | - | - | - | ↘ | ↓ | - | ↓ | ↘ | ↓ | ↓ | *na* | | | | | |
| | Bacteroidetes | Prevotellaceae | Prevotella | *na* | | | | | | *na* | | | | | | *na* | | | | | | *na* | | | | | | *na* | | | | | |
| | Firmicutes | Clostridiaceae | Faecalibacterium | *na* | | | | | | *na* | | | | | | *na* | | | | | | *na* | | | | | | *na* | | | | | |
| **Unclear** | Bacteroidetes | | | ↓ | ↘ | ↘ | ↘ | ↓ | ↓ | - | ↗ | - | - | - | - | ↘ | ↗ | - | - | ↘ | ↘ | - | - | - | - | ↓ | ↓ | - | ↗ | - | - | ↓ | ↓ |
| | Firmicutes | Erysipelotrichaceae | | ↗ | ↗ | ↘ | - | ↘ | ↑ | ↘ | ↘ | ↗ | ↗ | ↗ | ↗ | ↘ | - | ↘ | - | ↗ | - | ↘ | ↗ | ↓ | ↗ | ↑ | ↑ | ↘ | ↓ | ↘ | ↓ | ↑ | ↗ |
| | Firmicutes | Ruminococcaceae | | ↗ | ↗ | ↑ | ↗ | ↗ | - | ↗ | ↘ | - | ↘ | ↘ | ↘ | ↗ | ↘ | ↘ | ↘ | ↘ | - | ↑ | ↘ | ↑ | ↘ | ↗ | ↓ | - | - | ↗ | ↗ | ↓ | ↓ |

**Supplementary Fig. 1 Experimental design and in-life experiments**

a) Hierarchy graphs illustrating the complex experimental design. See details in Material and Methods. The white on black background numbers refer to the numbers of animals we were left with at the end of the in-life phase. b) Graph indicating which treatments were given or tests were performed during the in-life phase. Gross motor functions were tested weekly, while the adhesive removal test was only performed twice, at start and end of the in-life phase. Faeces were checked weekly and afterwards six time points were chosen for analysis. Mice were gavaged weekly starting at week 2 for a total 8 times and their food was changed every second week.

**Supplementary Fig. 2 Alpha and beta diversity assessment for the different challenges**

a) Boxplots for alpha diversity of the different variables/challenges. Genotype and Diet challenges were overall features altering alpha diversity. No significant changes were detected due to the Gavage challenge. Stats: Kruskal-Wallis, corrected for FDR. b) Non-metric multi-dimensional scaling (NMDS) plots for beta diversity of the different variables/challenges. Here Diet and Gavage appear to lead to changes in beta diversity. However, one has to take into account that the PBS gavaged mice were exclusively FD challenged. The dissimilarities were genotype independent. Stats: Adonis.

**Supplementary Fig. 3 Firmicutes to Bacteriodetes ratios increase under fibre deprivation**

Boxplots illustrating the Firmicutes to Bacteriodetes ratios for each tested time point between FR and FD challenged mice. Both genotypes show significant increases in ratios at 9 weeks. Thy1-Syn14 mice do already show significant differences at week 6.

Stats: Kruskal-Wallis, corrected for FDR

FD, fibre deprived; FR, fibre rich (normal chow); WT, wild-type littermates; TG, Thy1-Syn14

**Supplementary Fig. 4 Differential taxa abundance over time is diet regulated**

Complex heatmap visualizing the effect of different challenges on the relative abundance changes over time. The taxa were subdivided into three different groups according to their relative abundances: High, at least for one time point there was an average relative abundance of 10% or higher for at least one treatment group; Mid, at least for one time point there was an average relative abundance between at least 1 and maximum 10% for at least one treatment group; Low, less then 1%. All data was scaled and centred per row. Main effect is seen between FR and FD challenges. For some taxa we observe also difference between genotypes.

WT, wild-type littermates; TG, Thy1-Syn14; FD, fibre deprived; FR, fibre rich; PBS, phosphate buffered saline solution; ΔEC, curli-KO E.coli; EC, wild-type curli expressing E.coli

**Supplementary Fig. 5 Relative abundance of *E. coli – Shigella* from 16S rRNA amplicon sequencing**

Dotplots illustrating the relative abundance for Escherichia coli – Shigella. Tehse sequences were barely detected in twith the 16S rRNA amplicon sequencing approach.

**Supplementary Fig. 6 Endotoxin levels show great variation in between challenge groups**

Boxplots illustrating plasma endotoxin levels facetted by genotype. There are no significant differences between the different groups. We saw a notable inner-group variation. The endotoxin plasma are comparable to those seen in high fat diets (Kim et al. 2012).

*Kim, Kyung-Ah, Wan Gu, In-Ah Lee, Eun-Ha Joh, and Dong-Hyun Kim. 2012. 'High Fat Diet-Induced Gut Microbiota Exacerbates Inflammation and Obesity in Mice via the TLR4 Signaling Pathway'. PLoS ONE 7 (10): e47713. https://doi.org/10.1371/journal.pone.0047713.*

**Supplementary Fig. 7 Alpha diversity and mucus thickness correlate between diet challenges but in opposite directions**

Scatterplots of Spearman rank tests comparing alpha diversity (y-axis) and mucus thickness (x-axis) in both diet challenges separately. There are no significant correlations between microbial diversity and mucus thickness in neither diet groups. However, there are strong correlative trends in both diet groups. Interestingly, these trends are of opposite direction. Stats: Spearman rank test.

FD, fibre deprived; FR, fibre rich (normal chow)

**Supplementary Fig. 8 Challenges dependant pS129-αSyn positive accumulations in both genotypes**

Representative high magnitude (40x, Scale bar: 25μm) microscopic images of every group. Here we saw that there are pS129-αSyn+ accumulations in both genotypes. We did observe differences in particle size depending on the treatment combination. The greatest particles have been observed in FD EC challenges mice and more so in TG mice.

WT, wild-type littermates; TG, Thy1-Syn14; FD, fibre deprived; FR, fibre rich; PBS, phosphate buffered saline solution; ΔEC, curli-KO E.coli; EC, wild-type curli expressing E.coli

**Supplementary Fig. 9 Gross motor functions are transgene driven**

a,b) Longitudinal gross motor function monitoring visualized in line plots indicating the median value changes for the different treatment groups. Alpha-synuclein overexpression drives motor deficits in transgenic animals. a) Hind limb clasping scores increase within all transgenic groups (dotted lines) independent of their treatment. Differences between WT and TG animals were significant from baseline to week 9. Stats: Mann-Whitney U, corrected for FDR; ****, $p < 0.0001$. b) Grip strength results confirm that there is only a difference between the genotype and there are no observable difference due to any kind of treatment. Differences between WT and TG animals were significant from baseline to week 9. Stats: Mann-Whitney U, corrected for FDR; ****, $p < 0.0001$.

WT, wild-type littermates; TG, Thy1-Syn14; FD, fibre deprived; FR, fibre rich; PBS, phosphate buffered saline solution; ΔEC, curli-KO E.coli; EC, wild-type curli expressing E.coli

**Supplementary Fig. 10 Sensory abilities differ only between genotypes and do not exacerbate after diet and/or curli challenges**

Boxplots illustrating the latency for touch in the different treatment groups (x-axis) for WT (red) and TG (blue) animals after 9 weeks. The time of touch refers to the sensory ability of the mice. Here we saw a clear highly significance between genotype (Stats: Kruskal-Wallis, corrected for FDR; $p < 0.0001$). More specifically there are significant differences in the FR ΔEC, FR EC and FD EC between genotypes (Stats: Mann-Whitney U test, not corrected for FDR; *, $p < 0.05$; **, $p < 0.01$).

WT, wild-type littermates; TG, Thy1-Syn14; FD, fibre deprived; FR, fibre rich; PBS, phosphate buffered saline solution; ΔEC, curli-KO E.coli; EC, wild-type curli expressing E.coli

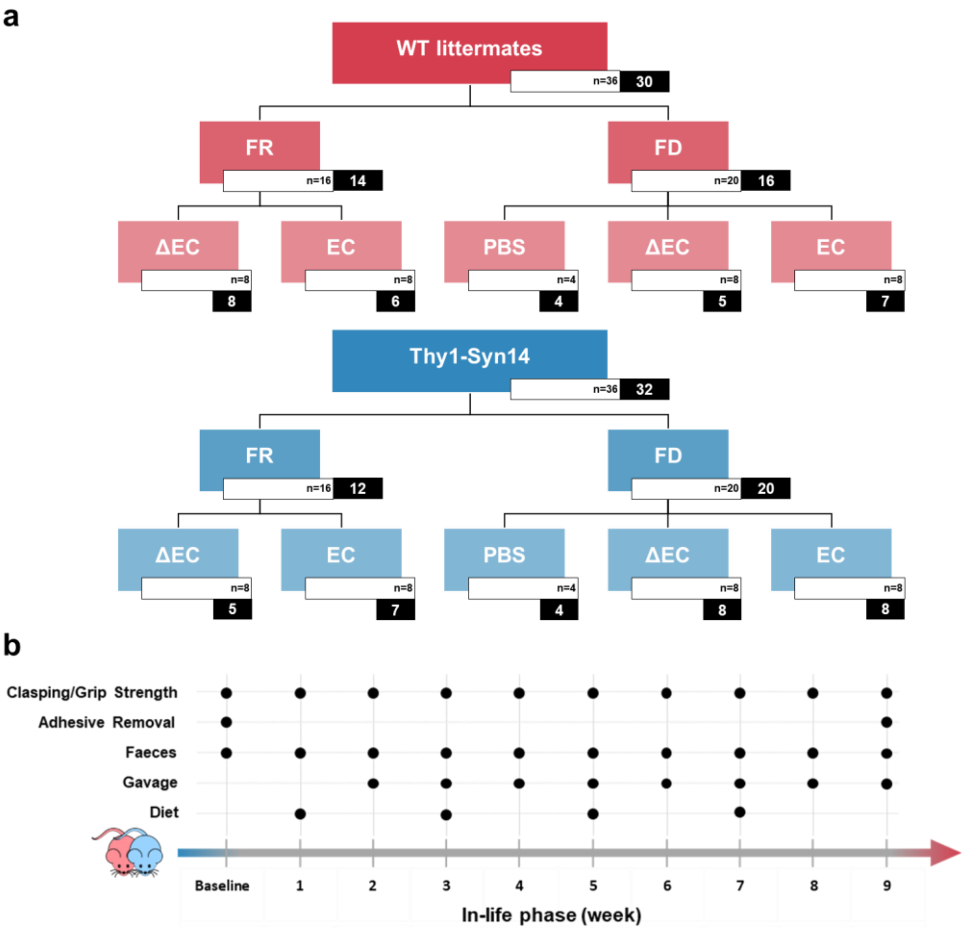


*Supplementary Fig. 1*

---

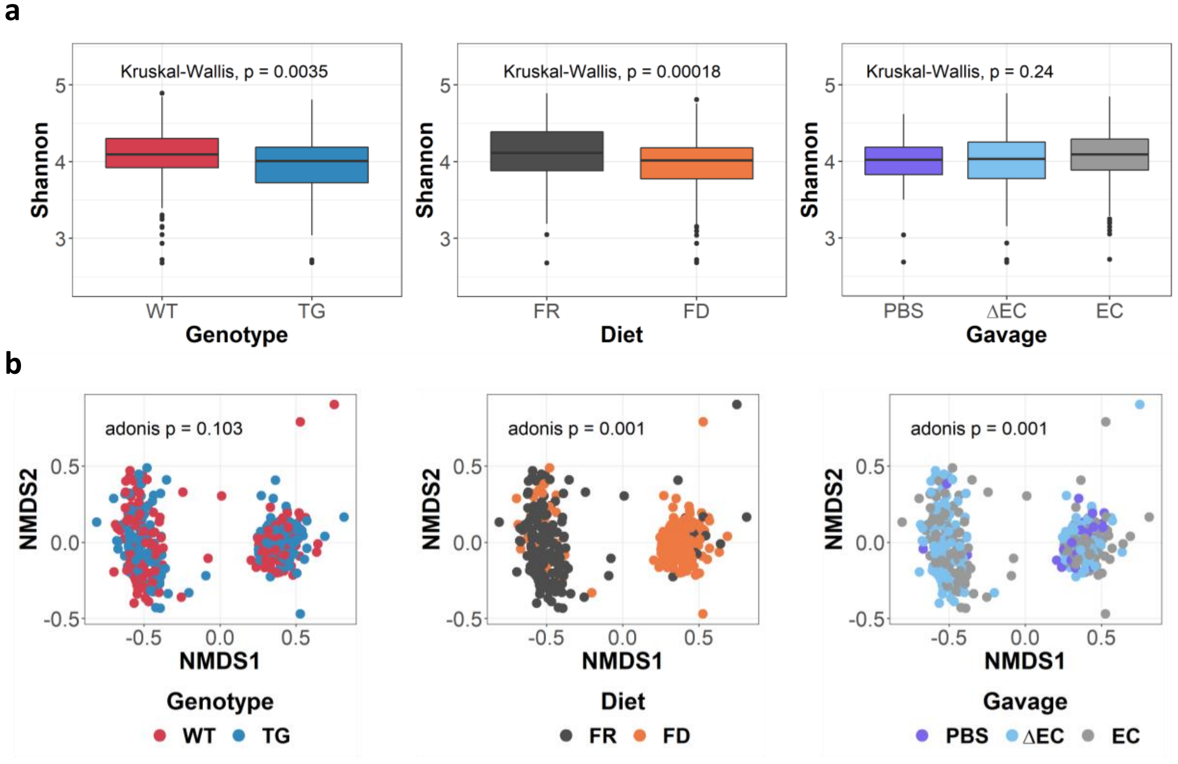


*Supplementary Fig. 2*

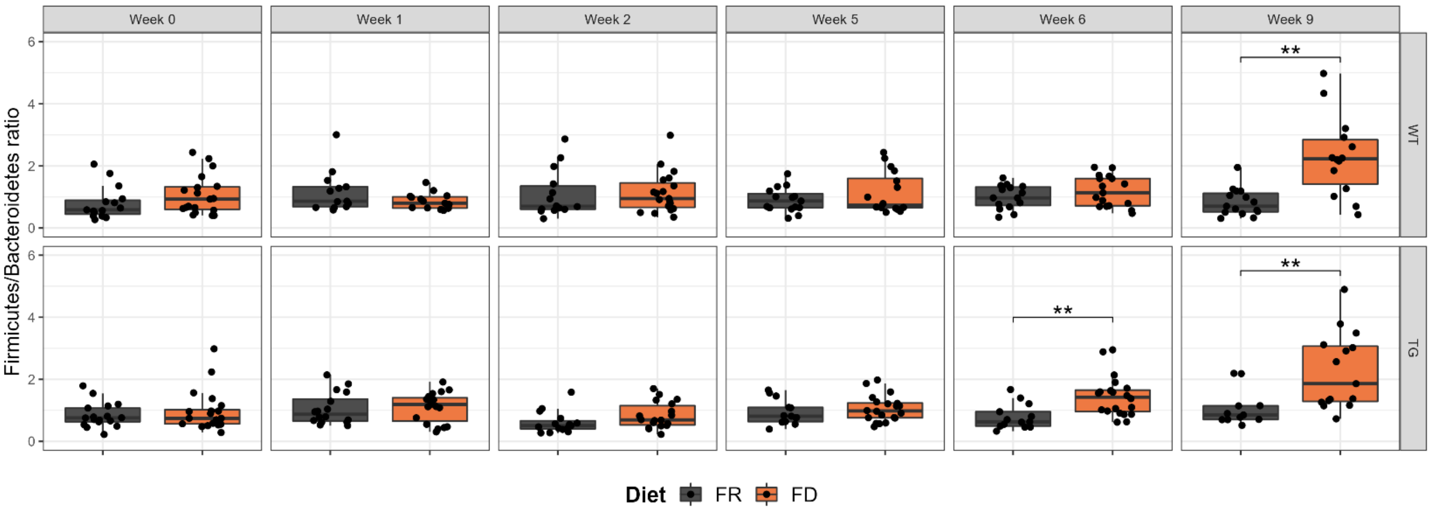


*Supplementary Fig. 3*

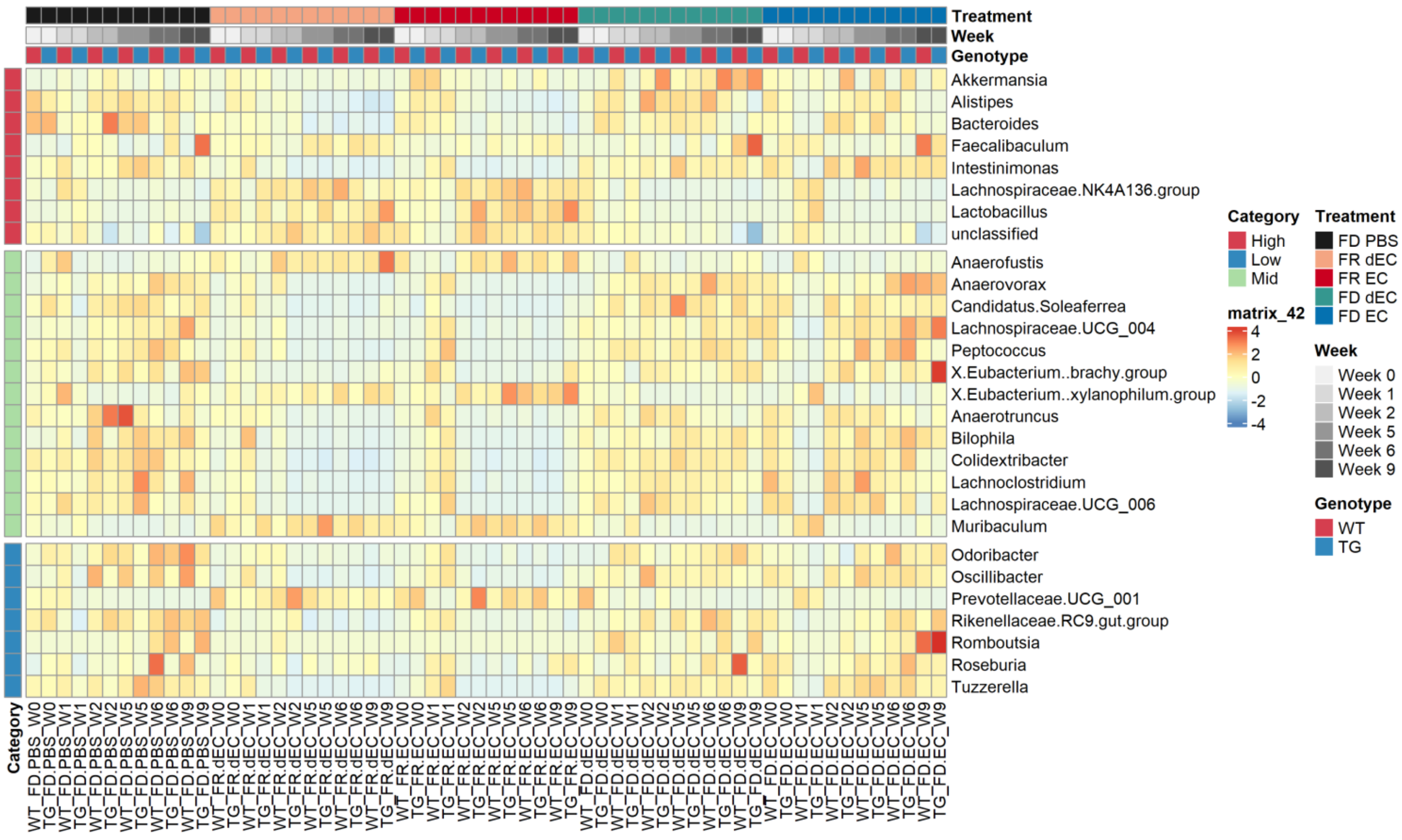


*Supplementary Fig. 4*

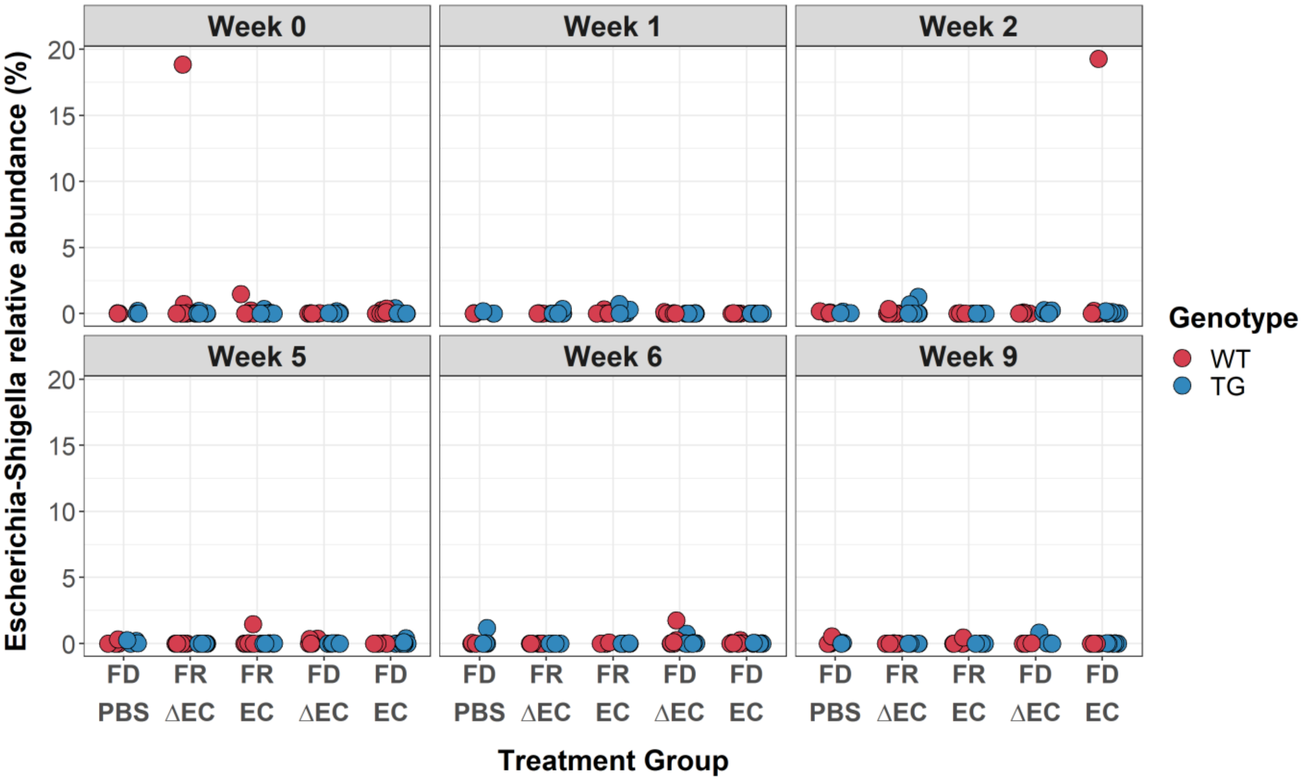


*Supplementary Fig. 5*

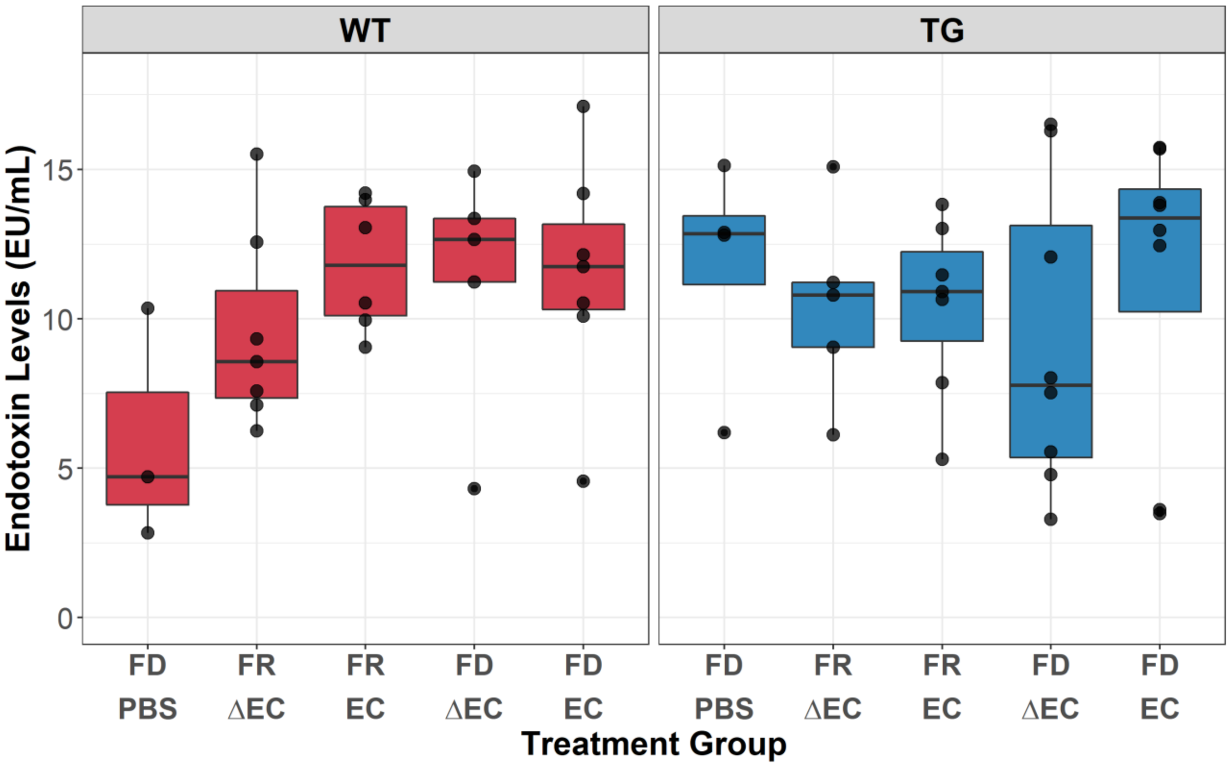


*Supplementary Fig. 6*

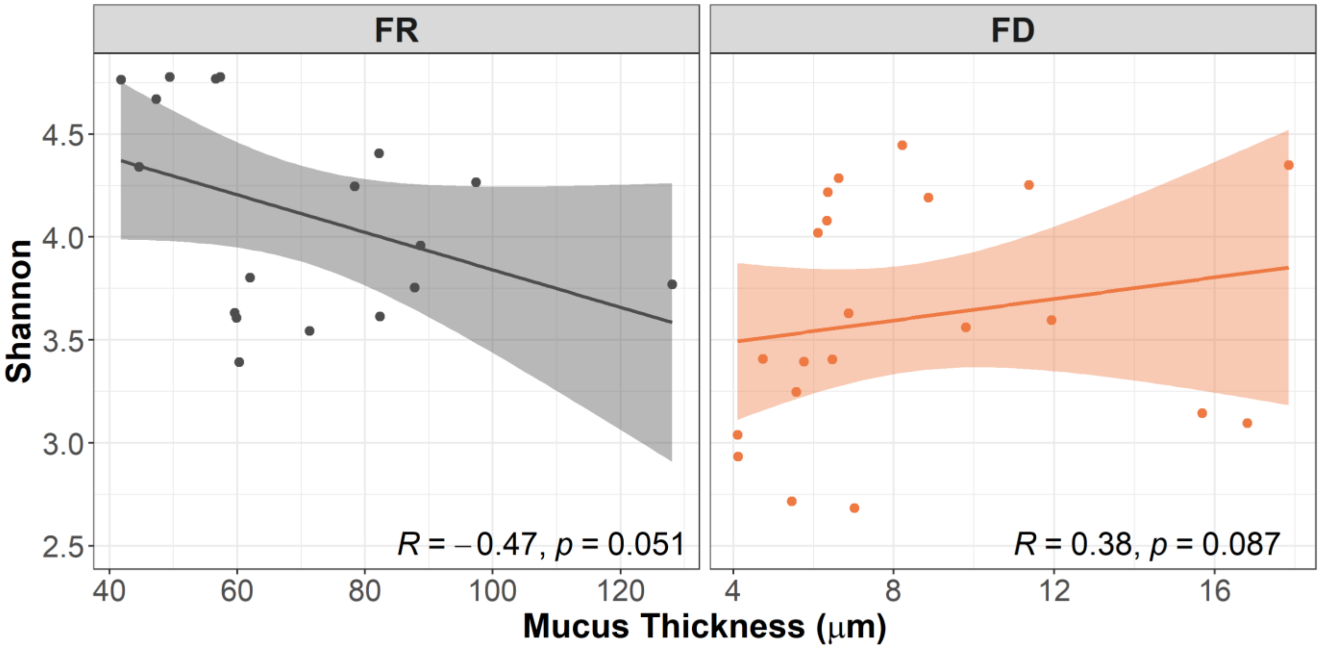


*Supplementary Fig. 7*

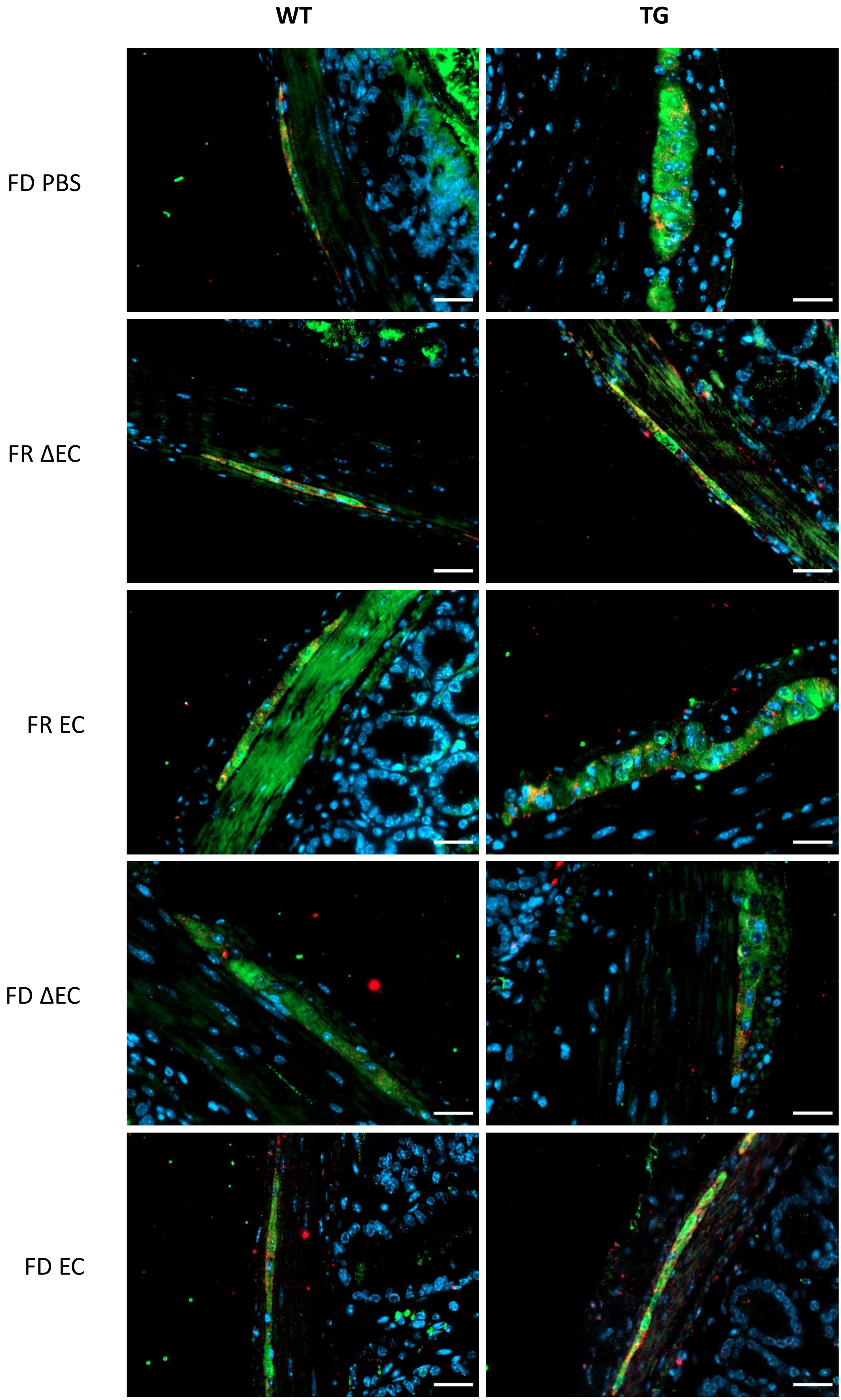


*Supplementary Fig. 8*

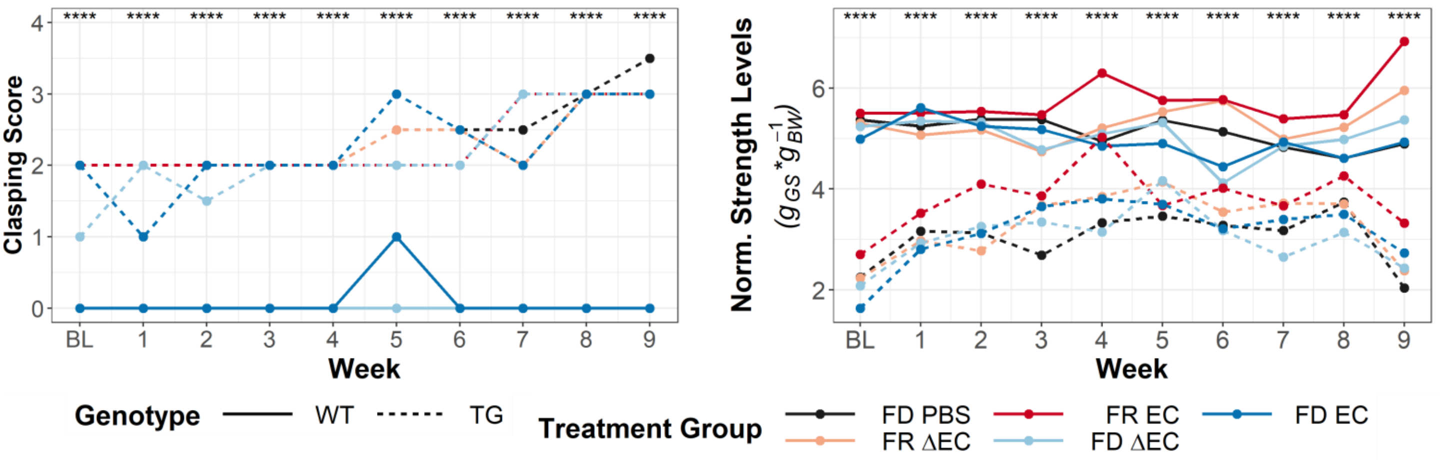


*Supplementary Fig. 9*

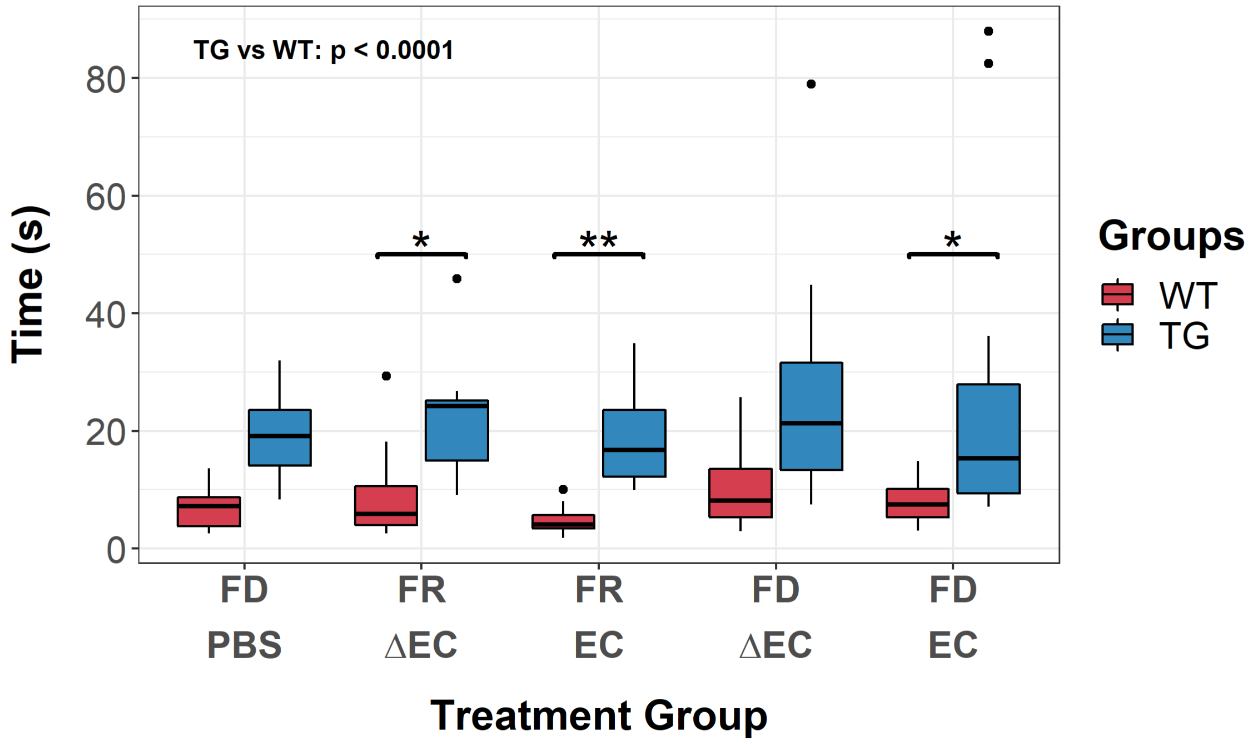


*Supplementary Fig. 10*
